# Characterising tandem repeat complexities across long-read sequencing platforms with TREAT and *otter*

**DOI:** 10.1101/2024.03.15.585288

**Authors:** Niccolo’ Tesi, Alex Salazar, Yaran Zhang, Sven van der Lee, Marc Hulsman, Lydian Knoop, Sanduni Wijesekera, Jana Krizova, Anne-Fleur Schneider, Maartje Pennings, Kristel Sleegers, Erik-Jan Kamsteeg, Marcel Reinders, Henne Holstege

## Abstract

Tandem repeats (TR) play important roles in genomic variation and disease risk in humans. Long-read sequencing allows for the accurate characterisation of TRs, however, the underlying bioinformatics perspectives remain challenging.

We present *otter* and TREAT: *otter* is a fast targeted local assembler, cross-compatible across different sequencing platforms. It is integrated in TREAT, an end-to-end workflow for TR characterisation, visualisation and analysis across multiple genomes.

In a comparison with existing tools based on long-read sequencing data from both Oxford Nanopore Technology (ONT, Simplex and Duplex) and PacBio (Sequel 2 and Revio), *otter* and TREAT achieved state-of-the-art genotyping and motif characterisation accuracy.

Applied to clinically relevant TRs, TREAT/*otter* significantly identified individuals with pathogenic TR expansions. When applied to a case-control setting, we significantly replicated previously reported associations of TRs with Alzheimer’s Disease, including those near or within *APOC1* (p=2.63×10-9), *SPI1* (p=6.5×10-3) and *ABCA7* (p=0.04) genes.

We finally used TREAT/*otter* to systematically evaluate potential biases when genotyping TRs using diverse ONT and PacBio long-read sequencing datasets. We showed that, in rare cases (0.06%), long-read sequencing suffers from coverage drops in TRs, including the disease-associated TRs in *ABCA7* and *RFC1* genes. Such coverage drops can lead to TR mis-genotyping, hampering the accurate characterisation of TR alleles.

Taken together, our tools can accurately genotype TR across different sequencing technologies and with minimal requirements, allowing end-to-end analysis and comparisons of TR in human genomes, with broad applications in research and clinical fields.

## 1. Introduction

Roughly 30% of the human genome consists of tandem repeats (TR) characterised by one or more repeat motifs that are defined by their consecutive repetition.^1^ This repetitive pattern often leads to DNA instability, facilitating not only expansions and contractions of the repeating motif sequence, but also allelic diversity within the sequence.^2,3^ Several definitions of TRs have been introduced based on the motif length and size variability, including microsatellites, minisatellites, and macrosatellites. Microsatellites (or short tandem repeats, STR) are the most abundant TRs in the human genome, are characterised by a repetitive motif of less than 6 base pairs (bp), and tend to cluster in non-coding regions of the genome.^4^ Minisatellites are characterised by a repetitive motif with a size ranging 7-100 bp, and they are highly enriched in the telomeric regions of the genome.^5^ Macrosatellites are characterised by larger tandem repeat units (>100 bp), and are enriched in the telomeric and centromeric portions of the genome.^6^

TRs can disrupt gene-expression regulation and contribute to over 40 neurological disorders.^1,7,8^ Pathogenic TR expansions, surpassing critical lengths, are linked to conditions like spinocerebellar ataxias, Huntington’s disease, Fragile-X syndrome, Amyotrophic lateral sclerosis (ALS), and Myotonic Dystrophy.^7–9^ For instance, Fragile-X syndrome results from a GGC repeat expansion in the *FMR1* gene, with affected individuals having up to 4,000 copies compared to less than 50 in healthy individuals.^10^ Similarly, ALS is caused by an intronic hexa-nucleotide repeat expansion (GCCCCG) in the *C9orf72* gene, exceeding a critical length of more than 200 copies.^11^ Beyond diseases-causing, TRs have been also identified as risk factor for complex human diseases: for example, the intronic TR in the *ABCA7* gene is associated with a 4.5-fold increased risk of Alzheimer’s Disease (AD) when the TR exceeds 5720 base pairs.^12,13^

Traditionally, the evaluation of TR lengths and sequences has been challenging. Conventional methods, such as repeat-primed polymerase chain reaction (RP-PCR) and Southern blot assays, are time-consuming and limited in detecting TRs within PCR-based boundaries. Short-read sequencing approaches offer an alternative, but their limited read lengths often fail to span repetitive regions effectively. Despite heuristic methods and statistical modelling,^14–19^ accurately assessing clinically relevant TRs remains difficult. The advent of long-read sequencing, particularly with PacBio’s High Fidelity (HiFi) and Oxford Nanopore Technology’s (ONT) Duplex technology (10-20kb on average, >99% accuracy),^20,21^ has significantly improved TR evaluation by providing long and accurate sequencing fragments.

Characterising TRs with long-read sequencing technology currently has two main limitations. First, there is the need to characterise TRs across different (long-read) sequencing technologies and data-types.^22–24^ This is critically important given the growing long-read sequencing initiatives aiming to comprehensively assess TRs in large genomic datasets,^25^ spanning both population-wide and clinical contexts. For example, some existing tools are constrained by predefined TR databases, hindering the identification of new TR features such as novel motif sequences;^26^ other tools are technology and data-type-dependent,^22^ or do not produce generalizable multi-sample outputs.^23,24^

Second, there is a lack of comprehensive studies that have investigated potential biases when sequencing TRs. For example, DNA methylation has been previously shown to impact base-calling accuracy in long-read sequencing data.^27–29^ Similarly, the formation of secondary structures due to TRs could impact enzyme efficiency (*e.g.* polymerase or nanopores),^30^ potentially reducing read-quality and sequencing throughput in current long-read sequencing technologies. Furthermore, some technologies require the alignment of noisy reads to generate high quality consensus sequences, which might be more difficult in case of repetitive regions. These problems may impact genotyping accuracy and lead to incorrect assessments of allele-sequences, including disease-associated TRs in patients.

Here, we present TREAT (Tandem REpeat Annotation Toolkit), a unified workflow for characterising TRs across multiple genomes, cross-compatible with diverse long-read technologies and data-types (*e.g.* read-alignments and *de novo* assemblies). TREAT employs a novel generic targeted local assembler, *otter*, that can adapt to different sequencing chemistries to accurately characterise TRs. We benchmarked TREAT and *otter* with currently available tools for TR genotyping (PacBio’s TRGT and LongTR)^22,31^ in terms of genotyping accuracy, motif identification, and running performances. We then showcase TREAT and *otter* applicability in a population-, clinical-, and case-control setting. Finally, we performed a systematic analysis of ∼864K genome-wide TRs in CHM13 reference genome to evaluate sporadic coverage drops that can affect TR genotyping accuracy. We did so using the well-characterised HG002 genome based on long-read sequencing data from ONT (Duplex and Simplex), HiFi and non-HiFi data from PacBio’s Revio and Sequel 2 instruments.

## 2. Results

### 2.1 Cross-compatible workflow for characterising tandem repeats with *otter* and TREAT

We present *otter* and TREAT, two bioinformatic tools that enable tandem repeat (TR) characterisation across different long-read sequencing technologies and data-types with minimal input requirements (*Figure 1*). *Otter* is a stand-alone generic targeted local assembler for long-read sequencing data, which automatically adapts to sequencing error-rates and coverage levels per target region. TREAT integrates *otter* to enable end-to-end unified workflow for *de novo* motif characterisation and downstream analysis, including TR visualisation, outlier-based and case-control comparisons (see *Methods*). Both tools require sequencing data aligned to a reference genome (.bam files), the reference genome used (.fasta file), and the coordinates of the regions of interest (chromosome, start and end positions encoded in a .bed file). TREAT/*otter* outputs a multi-sample gVCF (Genomic Variant Call Format) file reporting genotyped alleles, their size and relative repeat content (motif and number of copies), of each TR in each sample.

**Figure 1:**
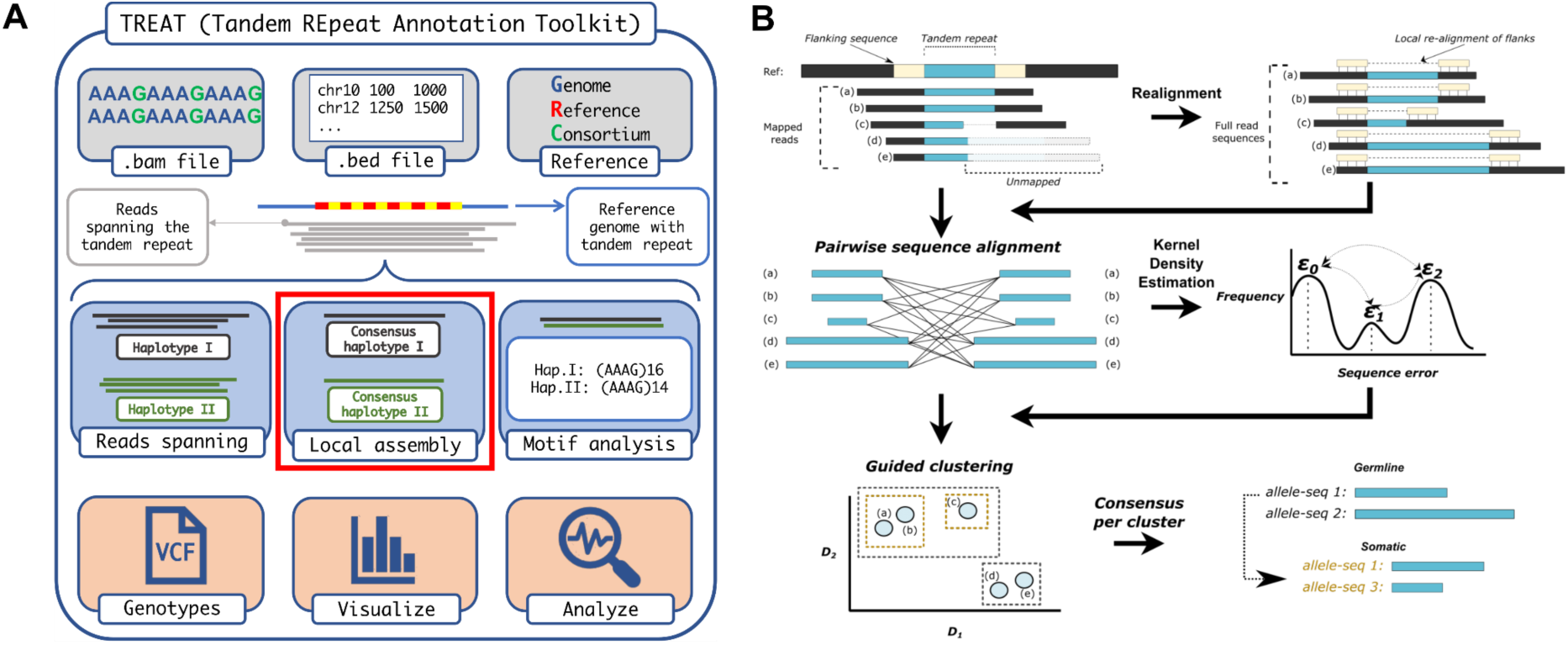
*Schematic workflow of TREAT and otter*. **A.** Shows TREAT workflow, highlighting the required inputs, the main features, and the main outputs of the tool. The red box highlights the main genotyping engine based on *otter.* **B.** Summarises the main algorithmic steps of *otter*, a novel targeted local assembler for long-read sequencing data.

*Otter* is written in C++ and the source code is freely available at https://github.com/holstegelab/otter.

TREAT is written in Python and R (for plots). The source code is freely available at https://github.com/holstegelab/treat along with example datasets, documentation, a dedicated Conda configuration file and a Docker image to ease the installation.

### 2.2 *Otter* and TREAT enable accurate characterisation of both PacBio and ONT long-read data

We benchmarked TREAT and *otter* with TRGT and LongTR,^22,31^ currently available tools to characterise TRs in long-read sequencing data. We compared: (*i*) genotyping accuracy, *i.e.* the accuracy of the predicted allele sequences for a TR, (*ii*) motif characterisation accuracy, and (*iii*) computational resources. We varied different long-read sequencing technologies (PacBio Sequel 2 and Revio, ONT Simplex and Duplex) as well as different coverages (5x, 10x, 15x, 20x, 25x, and 30x) of HG002.^32^ We focussed on a set of 161,382 TRs from PacBio’s repeat catalogue (see *Methods*). Predicted TR alleles were compared to the *expected* alleles based on the HG002 T2T assembly (see *Methods*).

In PacBio data, we found comparable genotyping accuracy between *otter* (TREAT genotyping engine) and TRGT, for both Sequel 2 and Revio datasets, although *otter* generated more accurate genotypes for larger TRs (*e.g.* >500bp), achieving average error-rates of 0.2-2.5%, compared to 0.6-3.8% of TRGT. Both methods were more accurate when increasing the coverage, although this was less pronounced for larger TRs (>500 bp). Notably, genotyping accuracy for both *otter* and TRGT was higher for PacBio’s Sequel 2 data in comparison with Revio data (*Figure 2A* and *Supplementary Results*). In ONT data, *otter* was generally more accurate than LongTR although differences for large TRs were less clear. For both tools, we observed better accuracies for Duplex data in comparison to Simplex data (*Figure 2B* and *Supplementary Results*). Altogether, our benchmark across all tools revealed that PacBio led to more accurate genotypes for TRs <500 bp, with PacBio and ONT having similar performances for TRs ranging 500-1000 bp, and ONT leading to more accurate genotypes for TRs >=1000 bp (*see Figure 2A-B and Supplementary Results*).

**Figure 2:**
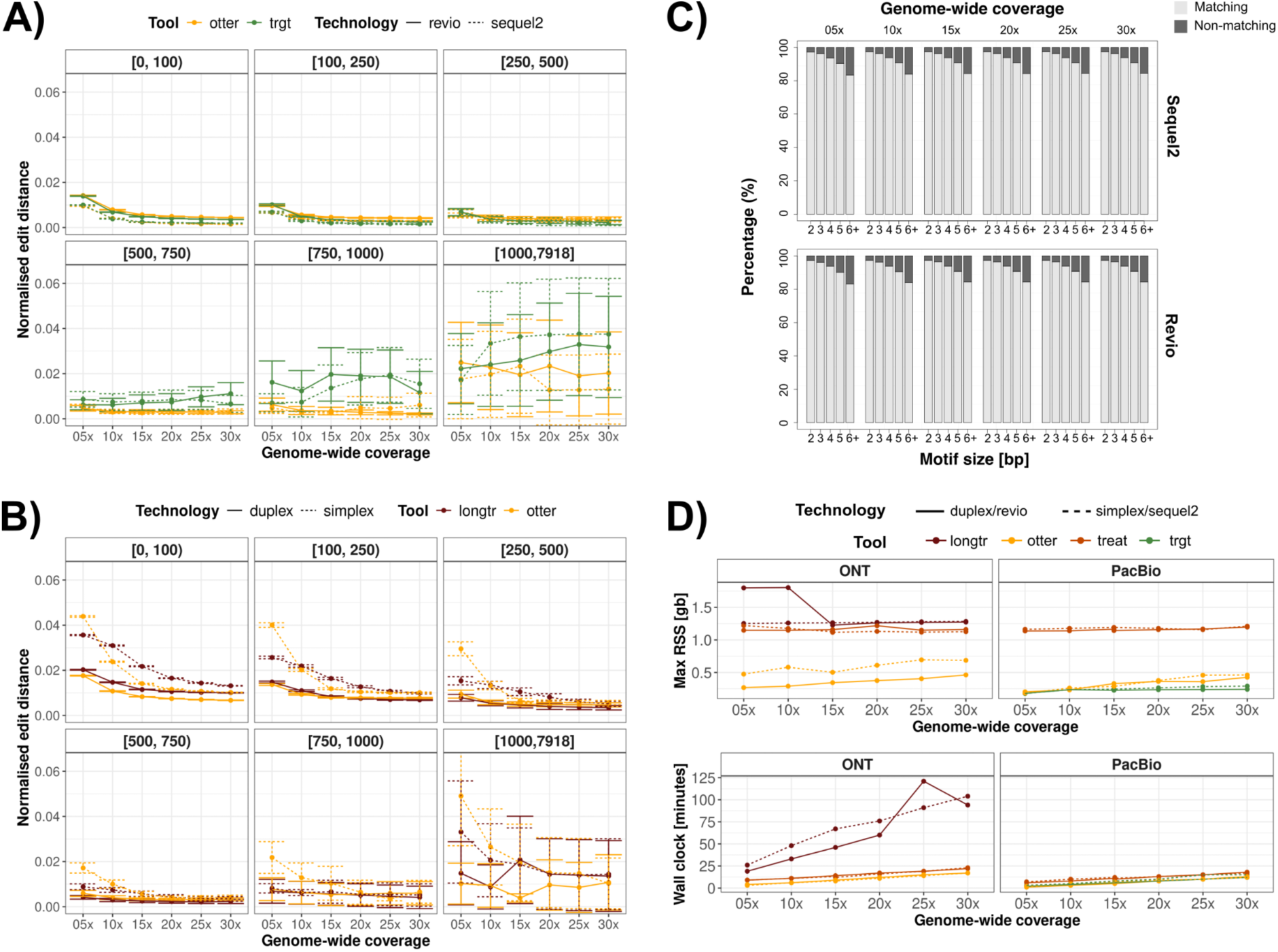
Benchmarking between TREAT/otter, TRGT, and LongTR. **A.** Genotyping accuracy of otter and TRGT on PacBio Sequel 2 and Revio data, stratified by TR size and sequencing depth. **B.** Genotyping accuracy of otter and LongTR on ONT Simplex and Duplex data, stratified by TR size and sequencing depth. **C.** Motif identification accuracy of TREAT and TRGT on PacBio Sequel 2 and Revio data, showing the overlap of matching motifs, stratified by motif size and sequencing depth. **D.** Memory usage and running time of otter, TREAT, TRGT, and LongTR, across technologies and sequencing coverages.

The above observations remain when using different distance metrics and partitioned by different TR-types. For example, we observed similar performances when using the raw edit distance and correlation between observed and expected allele sizes (*Figure S1* and *Supplementary Results*). Furthermore, we found that TRs characterised by dinucleotide repeat motifs were on average less accurate than TRs with longer motifs (*Figure S2*). The fraction of alleles perfectly genotyped (*i.e.* with an edit distance of 0), compared to expected alleles, increased with higher coverage across all technologies and tools (*Figure S3*), with Sequel 2 data having the largest fraction of alleles perfectly matched, and ONT Simplex having the least. In PacBio Sequel 2 and Revio data, TRGT generated a slightly higher fraction of perfectly matched alleles with respect to *otter* (max difference 2.8%). In ONT data, *otter* outperformed LongTR in all settings.

Similarly, TREAT, which makes use of TR-genotypes from *otter*, achieved similar motif characterisation accuracy relative to TRGT (*Figure 2C*). In the GRCh38 reference genome, the motifs of the 161K TRs were mostly dinucleotide (49%), followed by tetranucleotide (22%) and 16+ bp motifs (11%) (*Figure S4*). Because LongTR does not directly report the identified TR motifs, we compared TR motifs between TREAT and TRGT. On average, TREAT identified the same motifs as TRGT in 96% of cases (*Figure 2C*), and this did not change for different technologies (Sequel 2 or Revio) or different coverages. We observed a higher concordance in motif detection between tools for shorter motifs (*Figure 2C*). When looking at the motifs identified by TREAT on the GRCh38 reference genome, these matched known TR annotations in 91% of the cases.

Finally, we evaluated the computational performances of *otter* (stand-alone), TREAT (integrated workflow with *otter*), TRGT and LongTR. When using four threads, TRGT and *otter* had similar run-time performances, while both were slightly faster than the integrated workflow from TREAT (*Figure 2D*). On the other hand, for the ONT data, *otter* and TREAT were faster than LongTR. In terms of memory consumption, performances were comparable between TREAT and LongTR, while *otter* and TRGT used significantly less memory (*Figure 2D*). When evaluating the multithreading capabilities in TREAT, we saw that when increasing the number of threads to 6, 8, 10 and 12, the running times decreased by 1.3-, 1.5-, 1.6- and 1.8-fold (on average across the different technologies), compared to 4 CPU threads (*Figure S5*).

In addition to the high-quality HiFi data, PacBio can output non-HiFi data, *i.e.* reads that did not pass PacBio’s internal HiFi quality thresholds, and that constitute a significant fraction of all sequenced data (45% in HG002). We explored whether integrating both HiFi and non-HiFi data could improve *otter*’s capability to accurately characterise TR allele sequences. Because Revio uses a subset of these non-HiFi reads (those with at least 90% read quality) to improve throughput and accuracy via DeepConsensus,^33^ we performed this analysis only for Sequel 2 data. We found that non-HiFi data improved accuracy across all TR-lengths. Specifically, when integrating non-HiFi reads of at least 85-90% read quality, genotyping accuracy improved by nearly two-fold (*Figure S6*).

### 2.3 TREAT’s unified workflow enables diverse characterisations of tandem repeats

We applied TREAT’s unified workflow to characterise TRs in a population and clinical setting. First, we genotyped the set of 161K TRs in 47 genomes from the Human Pangenome Research Consortium (HPRC),^34^ for which PacBio HiFi data was available. We then extracted the top 20% most variable TRs (N=32,208, based on the coefficient of variation, see *Methods*), and performed a principal component analysis (PCA, *Figure 3A*) on the joint allele sizes (*i.e.* the sum of the maternal and paternal alleles). We found that PC1 explained 12% of the total variance and genetically represented the African-American axis, while PC2 explained 3.5% of variance and corresponded to the American-Asian axis. The explained variance was similar to that of a PCA including 40/47 matching samples and 30,544 random common (minor allele frequency >10%) Single Nucleotide Polymorphisms (SNPs) (PC1: 14%, PC2: 4%, *Figure S7*).

**Figure 3:**
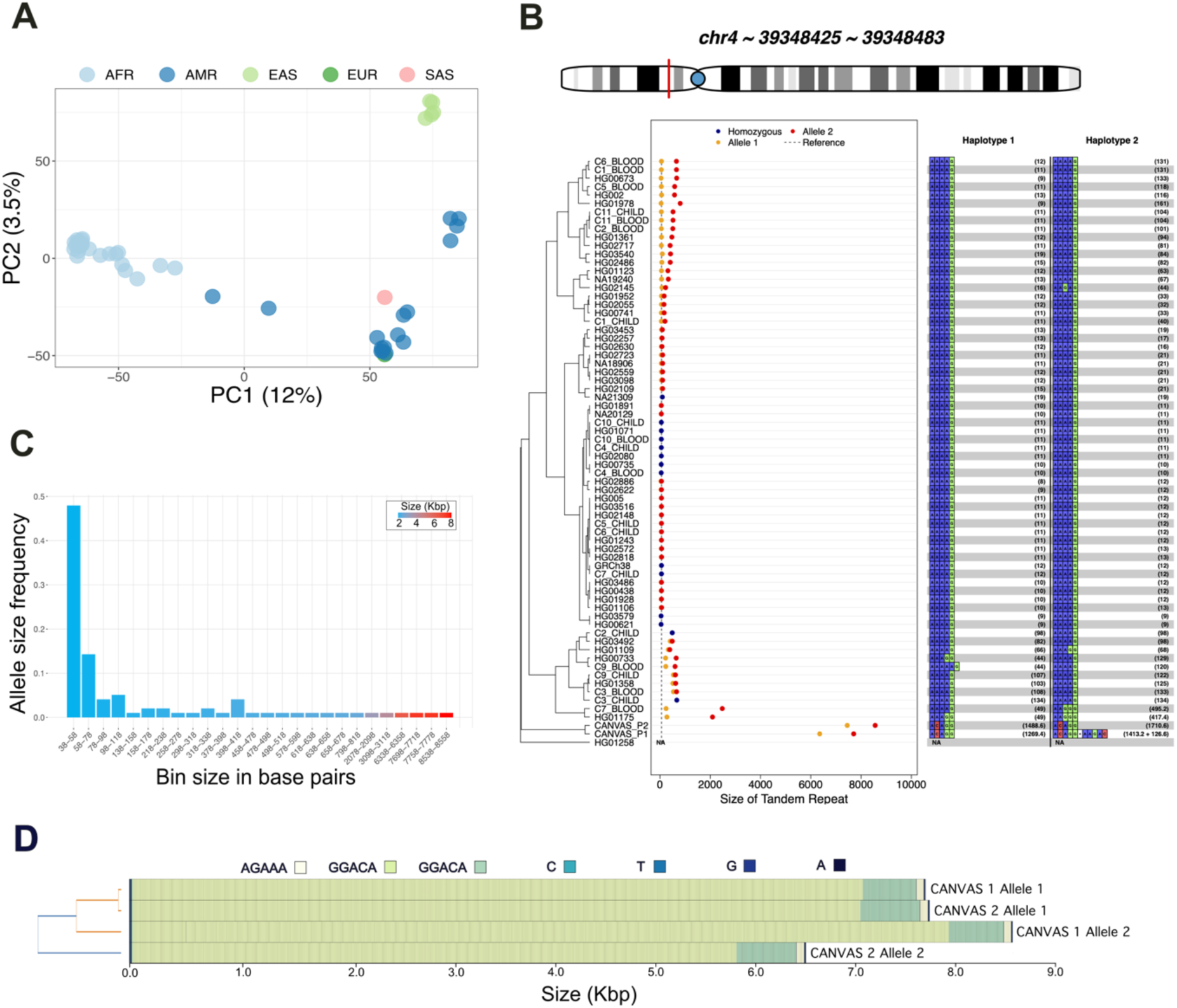
TREAT visualisation and analysis modules. **A.** The PCA of the ancestry-based analysis based on the 20% most variable TRs across 47 HPRC genomes. **B.** The main TR in the RFC1 gene. Y-axis: individuals, X-axis: TR size (in bp). Blue dots: smaller allele, orange dots: larger allele, red dots; homozygous genotypes. Dashed line: the allele in the reference genome GRCh38. The right side of the plot reports, for each sample and each allele, the motif and relative number of copies. The TR length of the two CANVAS patients were identified as significant outliers compared to the length-distribution of 47 samples from the HPRC. **C.** The distribution of allele sizes for the TR in RFC1 gene. **D.** Motif representation in CANVAS patients, as produced with MotifScope.^37^

We then used TREAT’s outlier analysis to detect and score extreme TR expansions or contractions of 35 clinically relevant TRs (*Table S1*) in 47 genomes from the HPRC, as well as two Dutch CANVAS patients and 10 parent-offspring duos (see *Methods*).^35,36^ The two CANVAS patients were previously characterised to harbour expansions in the intronic TR in *RFC1*.^35^ For all individuals, PacBio HiFi data were generated with Sequel 2 instrument. In total, we identified 30 instances where the TR length in certain samples were significantly different from the distribution of TR lengths across all 69 genomes. The most significant deviations were observed for the two CANVAS patients in the TR intronic of *RFC1* gene (*p*<2×10-16 for both patients, *Figure 3B-D*). The joint allele size for these samples was 78- and 89-fold higher than the median TR size across all 69 genomes. Significant TR expansions were also found in the TR in *ATXN8* gene (HG01123 sample, *p*<2×10-16, *Figure S8*), and in *DMD* gene (HG02622 sample, *p*=6.90×10-3, *Figure S9*). Interestingly, in the TR intronic of *RFC1* gene, we also observed a significant heterozygous expansion in one parent of the parent-offspring duos (p=1.7×10-3 and p=5.18×10-11, respectively for the short and long alleles, *Figure 3B*). Unexpectedly, the child reported a homozygous non-expanded genotype, suggesting a mis-assembly or an allele dropout.

Finally, we applied TREAT to characterise unique TRs that are present in CHM13 reference genome but absent in GRCh38 across the 47 HPRC genomes. We first curated a set of ∼864K genome-wide TRs in the CHM13 reference genome (see *Methods*). We evaluated genotyping accuracy by applying TREAT/*otter* to CHM13-aligned long-read datasets of HG002 (PacBio’s Revio and Sequel 2 as well as ONT’s Duplex and Simplex). We observed similar performances as those observed when using ∼161K TRs from GRCh38 (see *Figure S10* and *Supplementary Results*). These results showcase *otter* and TREAT’s ability to *de novo* characterise TRs across different reference genomes, and without prior knowledge of TR motif composition. Based on a CHM13-to-GRCH38 liftover procedure, we found 1017 unique TRs present in CHM13 and absent in GRCh38, 37% of which overlapped coding sequences (*Supplementary Methods* and *Table S2*). We used TREAT/*otter* to characterise these TRs across the 47 HPRC genomes and found a mean TR size of 129 bp (median=45 bp), mainly composed of trinucleotide motifs (42%), followed by homopolymers (26%), and 6+ nucleotide motifs (22%, *Figure S11*).

### 2.4 Tandem repeats may be sensitive to coverage dropouts in long-read sequencing

A closer investigation of PacBio long-read data revealed unexpected drops of coverage in clinical TRs, consequently leading to mis-genotyping of disease-associated TRs. One example is the CANVAS-associated intronic TR in *RFC1,* where the most common allele consists of an (AAAAG)11 motif, with a total size of ∼55bp. In CANVAS patients, the TR can range from 2-10 Kbp in total length (*Figure 3B-D* and *Supplementary Results*). In one parent-child duo, we found that the parent harboured an expanded heterozygous version of the TR: a shorter allele with a total length of 244 bp with the (AAAAG)50 motif; and a longer allele with a total length of 2.49 Kbp, composed primarily of the (AAGGG)490 motifs (*Figure 4A*). Long-read sequencing of brain tissue from the same individual (PacBio Sequel 2) confirmed these results, although the longer allele was further expanded by 180 bp (36 additional motif-copies), suggesting a somatic expansion in the brain relative to blood (*Figure 4A*). However, long-read data from the child yielded a homozygous allele-sequence of 63 bp with the (AAAAG)12 motif (*Figure 4A*). This was unexpected as at least one of the two allele-sequences from the parent should be inherited in the child. A closer analysis of HiFi long-read-pileup overwhelmingly supported this genotype. However, we observed an abnormal coverage drop in both the parent and child for this TR, which was alleviated when including non-HiFi data (*Supplementary Results*). After merging HiFi and non-HiFi data of the child, TREAT/*otter* correctly assembled the expanded allele-sequence at 2.65 Kbp in size with (AAGGG)>374. Penta-repeat primed PCR (RP-PCR) confirmed that both parent and child harboured repeat expansions separately composed of the (AAAAG) and (AAGGG) motifs (*Figure S12*). Therefore, HiFi data alone failed to capture this expanded allele-sequence, which was recoverable when including the non-HiFi data.

**Figure 4:**
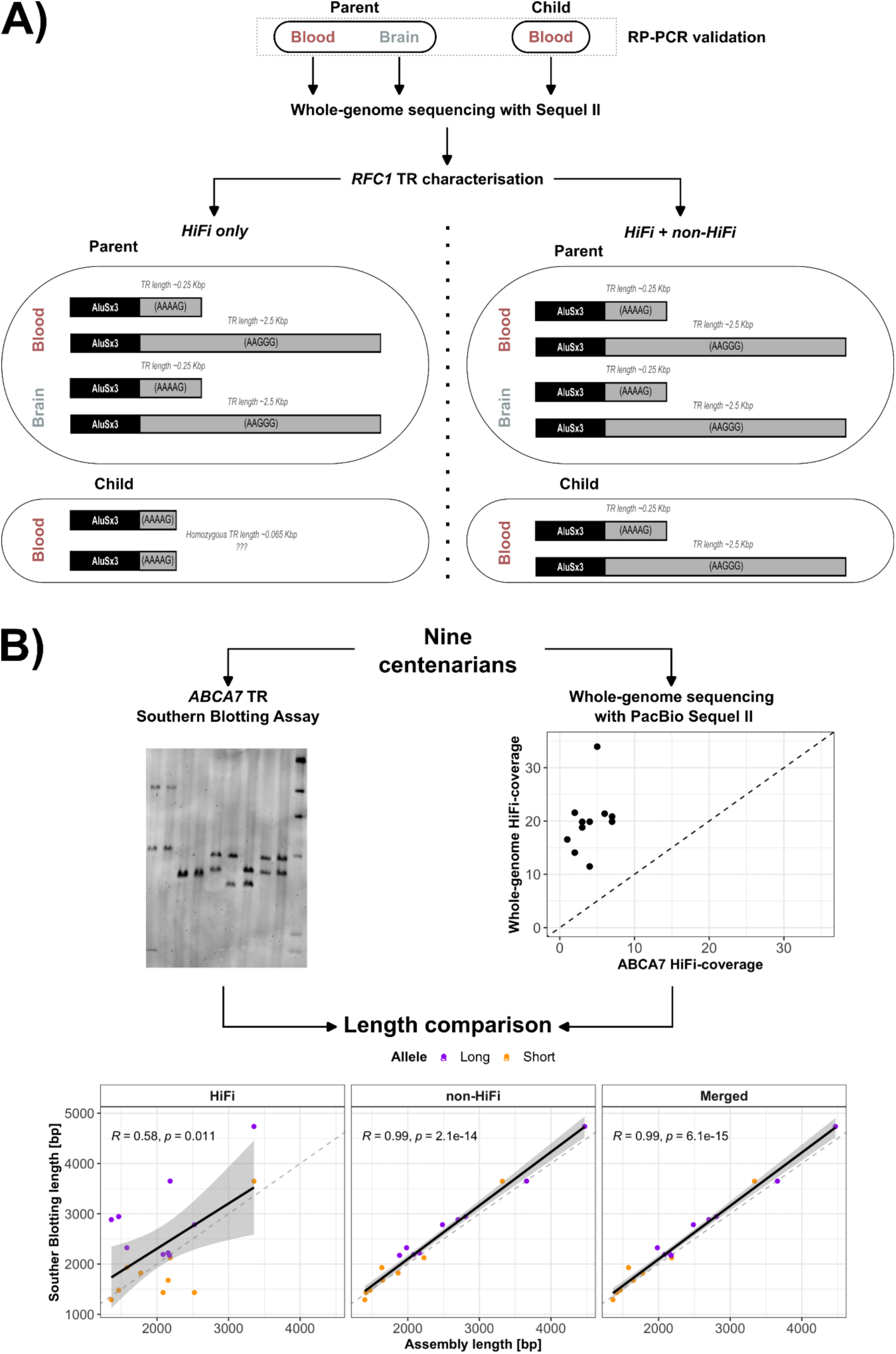
Coverage drops in TR in RFC1 and ABCA7 genes, associated with CANVAS and Alzheimer’s Disease. **A.** Shows the genotyped TR alleles and relative motif characterisation in a parent-child duo using only HiFi data, and HiFi + non-HiFi data. Long-read data from the brain of the parent were also available. Adding non-HiFi data rescued the missing allele in the child. **B.** Shows the comparison between experimentally validated alleles in the TR intronic of ABCA7 gene, and genotyped alleles based on HiFi data alone, and HiFi + non-HiFi. Experimental validation of TR alleles was performed with Southern Blot assay, and was available for 9 individuals for which long-read data was also available. When adding non-HiFi data, we could recover the expanded alleles in the child that were missed by HiFi data alone.

We observed similar situations of abnormal coverage drops in PacBio data in a separate intronic TR in *ABCA7*, previously associated with Alzheimer’s disease (AD). We experimentally validated the lengths of this TR using Southern Blotting in a subset of nine centenarians for which long-read sequencing was performed (*Figure S13* and *Supplementary Methods*). The local HiFi coverage for these individuals ranged 1-7x (*Figure 4B* and *Supplementary Results*). The correlation between experimentally validated alleles and HiFi-based alleles was 0.58 (Pearson correlation, *Figure 4B*). However, the inclusion of non-HiFi data increased read-support by four-fold to an average coverage of 22x. As a result, the correlation with experimentally validated allele sizes increased to 0.99 (*Figure 4B*). These results highlight standing challenges of characterising TRs with long-read sequencing data, and suggest systematic biases of long-read sequencing in certain genomic regions.

The above observations motivated us to systematically characterise genome-wide coverage drops of TRs across long-read sequencing technologies. We did this by investigating coverage drops in the curated set of ∼864K genome-wide TRs in the CHM13 reference genome, using both PacBio and ONT long-read datasets of HG002 at ∼38x coverage (see *Supplemental Results and Methods*). The average TR-length in this curated set was 93 bp, with motifs being mostly 16+bp motifs (23%), followed by dinucleotide (18%), tetranucleotide (14%), and homopolymers (13%, *Figure S4)*. For each TR, we defined the *coverage ratio* by dividing the local TR coverage *vs.* global genome-wide coverage. We found the average *coverage ratio* to be 1.01, 1.02, 0.99 and 1.03, respectively for Sequel 2, Revio, ONT Simplex and Duplex technologies. This indicated generally no unexpected coverage-drops in TRs (*Figure S14A*). However, 486 (0.06%) unique TRs had ratios below 0.25 (*i.e.* a four-fold lower coverage than expected based on the global average coverage), of which 454 (93%) were present in the HG002 T2T reference assembly (*Table S3*). The majority of the low-coverage TRs (294/454, 65%) overlapped gene annotations, potentially leading to mis-genotyping that may impact biological interpretation. Furthermore, we observed that some of these TRs were within 5 Kbp of each other, suggesting that coverage drops can extend across multi-Kbp regions. Overall, we observe significantly more low-coverage TRs in PacBio datasets compared to ONT (OR=9.4, p-value<2×10-16, Fisher’s exact test), with N=437 TRs (89%) being specific to PacBio datasets. Moreover, 22% of these TRs (N=98) had low coverage in both Sequel 2 and Revio datasets, suggesting potential systematic challenges in both technologies (*Figure S14B-G*). This included the intronic TR in *ABCA7*, previously associated with Alzheimer’s disease. Interestingly, the average number of non-HiFi reads in these TRs was 10, indicating that although reads were generated for these TRs, most were flagged as low-quality during HiFi data generation.

Within the ONT datasets, we observe significantly more low-coverage TRs in the Duplex dataset relative to the Simplex dataset (OR=2.6, p-value=1.76×10-3, Fisher’s exact test).

We characterised the sequences of all low-coverage TRs to investigate potential characteristic features. When comparing the 454 low-coverage TRs with the remaining of ∼864K genome-wide TRs, we found that low-coverage TRs were longer (p-value = 8.68e-14; 493 bp longer on average) and harboured higher GC-content (p-value = 2.28e-50; 17.4% higher on average). A comparison of dinucleotide content revealed that AG, CC, CG, CT, and GG dinucleotides were significantly enriched in the low-coverage TRs (*Figure S14H-I*). Moreover, we found that G-quadruplex DNA secondary structures (G4s) were more likely to occur in low-coverage TRs (p-value=2.48e-45; 3.76% higher, *Figure S14H* and *Supplementary Methods*).

### 2.5 Comparing tandem repeats across multiple genomes in a case-control setting

With the acquired knowledge about possible allele dropouts in TRs, we used TREAT/*otter* in a case-control setting to replicate the association of four TRs that were previously shown to associate with Alzheimer’s Disease (AD) risk (*Table 1, Table S4*). We did so by using a set of 246 AD patients (mean age = 67.9±9.8, 70% females) and N=248 cognitively healthy centenarians (mean age = 101.2±2.5, 70% females) that were sequenced with PacBio Sequel 2 instrument (*Methods* and *Figure S15*).^36^ Across all 494 genomes, we observed a median coverage (HiFi data) of 14, 15, 14, and 4, respectively for the TRs in *APOC1*, *SPI1*, *FERMT2*, and *ABCA7* (*Figure 5A*). The combined allele size (*i.e.* the sum of the maternal and paternal alleles) of the TR nearby *APOC1* (chr19:44921096-44921134) was significantly expanded in AD patients compared to cognitively healthy centenarians (beta=0.38, p=2.63×10-9, *Figure 5B* and *Table 1*). In contrast, the short allele of the TR within *SPI1* gene was significantly contracted in AD patients compared to cognitively healthy centenarians (beta=-0.03, p=6.5×10-3, *Figure 5B* and *Table 1*). The direction of effect of these TRs was in line with the original studies.^38,39^ We could not replicate the association of the TR within *FERMT2* (beta=0.01, p=0.27, short allele) (*Figure 5B* and *Table 1*).

**Figure 5:**
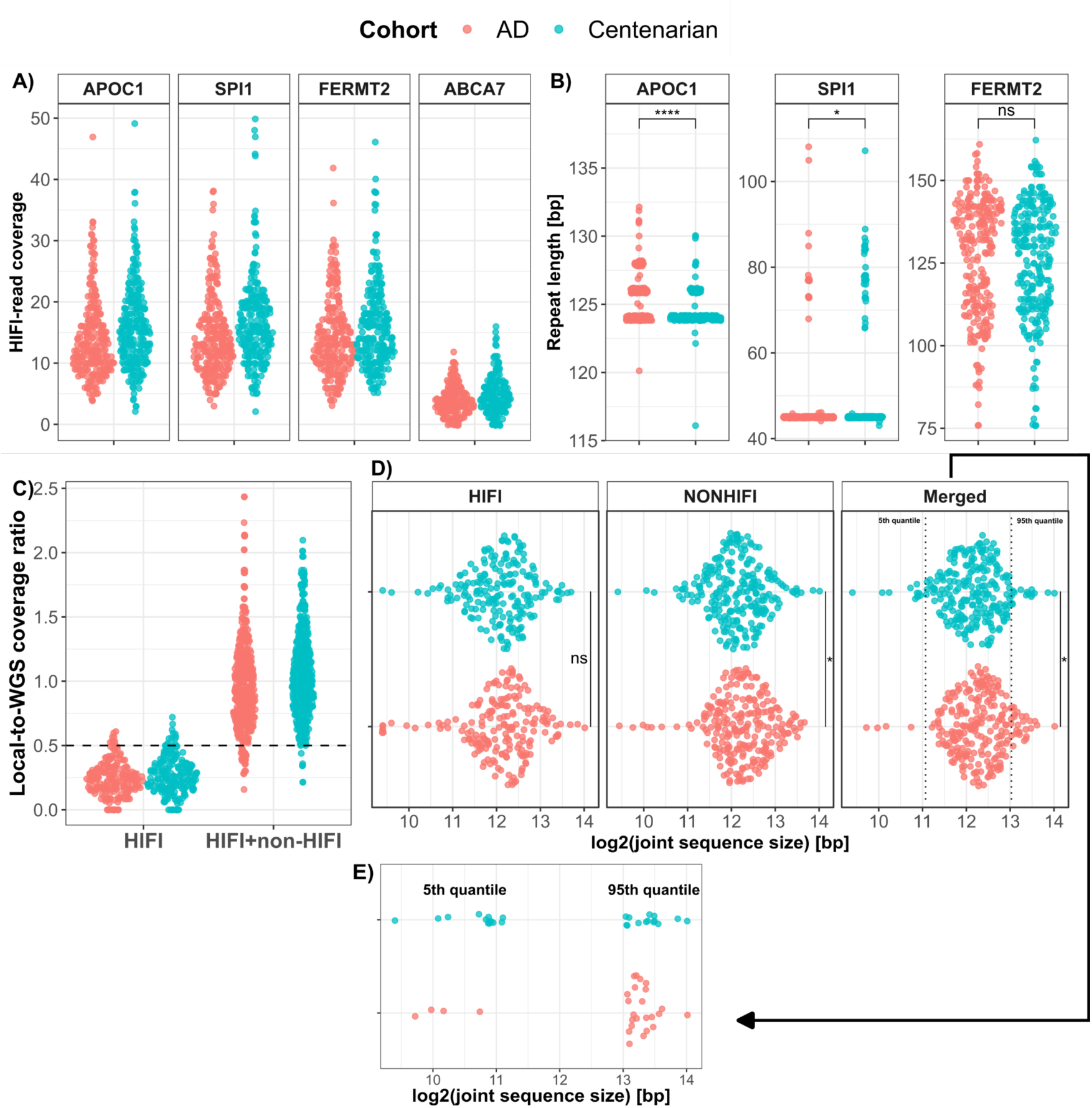
Replication of the association with AD of TRs in APOC1, SPI1, FERMT2 and ABCA7. **A.** The coverage distribution of the four TRs in AD patients and cognitively healthy centenarians. **B.** The TR size difference between AD patients and cognitively healthy centenarians in APOC1, SPI1 and FERMT2. For the associations, we used logistic regression models using the TR size as predictor for AD case-control status. **C.** HiFi and combined HiFi + non-HiFi coverage distribution of the TR intronic of ABCA7 gene. **D.** Comparison of the joint allele size of ABCA7 TR between AD cases and cognitively healthy centenarians, respectively using HiFi data, non-HiFi data, and the merged dataset of HiFi and non-HiFi. **E.** Number of AD cases and cognitively healthy centenarians in the lower 5^th^ quantile and upper 95^th^ quantile. Quantiles were defined based on the distribution of the joint TR-allele size in the centenarians. We tested for the differential enrichment of AD and centenarians in each quantile with Fisher’s exact tests.

**Table 1:** Replication of TR previously associated with Alzheimer’s Disease (AD)

|  | **TRs previously associated with AD** |  |  |  |
| --- | --- | --- | --- | --- |
| **Region** | chr19:44921096-44921134 | chr11:47775208-47775243 | chr19:1049436-1050066 | chr14:52832909-52832938 |
| **Gene** | APOC1 | SPI1 | ABCA7 | FERMT2 |
| **Best model** | Joint alleles | Short alleles | Joint alleles | Short alleles |
| **Beta (OR)** | 0.38 (1.46) | -0.03 (0.97) | 8.63x10<sup>-5</sup> (1.01) | 0.01 (1.01) |
| **P-value** | 2.6x10<sup>-9</sup> | 6.5x10<sup>-3</sup> | 0.041 | 0.27 |
| **Original study** | 38014121 | 37745545 | 29589097 | 37745545 |
| **Original OR** | NA | -0.01 (0.99) | 4.5 | 0.01 (1.01) |
| **Original model** | Longer allele | Joint alleles | Individuals with alleles >5720 bp | Joint alleles |
| **Original method** | Logistic regression | Mixed linear models | Fisher's exact | Mixed linear models |
| **Original p-value** | 4.3x10<sup>-10</sup> | NA | 0.008 | NA |
| **Original samples** | 1489 AD vs. 1492 controls | 6328 AD vs. 6580 controls | 275 AD vs. 177 controls | 6328 AD vs. 6580 controls |
| **Data type** | Short read sequencing | Short read sequencing | Southern blot | Short read sequencing |

Region: genomic coordinates of the TR with respect to GRCh38; Gene: the closest gene as reported in the original publications; Best model: model that yielded the most significant association, in our comparison: Short allele, Long allele or Joint alleles size; Beta (OR): effect size and relative Odds Ratio with respect to AD: an increased TR size leads to increased AD risk for positive estimates; P-value: p-value of association. We used logistic regression models using TR size (short allele, long allele and combined allele size) as predictor for AD case-control status, using 246 AD patients (cases) and 248 cognitively healthy centenarians (controls); Original study: the Pubmed ID of the original study; Original OR: the odds ratio as reported in the original study; Original model: model used for association in the original study; Original method: method used for association in the original study; Original p-value: the
*p-value reported in the original study; Original samples: the number of AD cases and controls used in* *the original study; Data type: the data on which the association were identified.*

For the intronic TR in *ABCA7*, we found significant expansions in AD cases after integrating non-HiFi data (beta=8.63×10-5, p=0.04, joint allele size, *Figure 5C-D*). We note that 22 samples were omitted due to reduced coverage levels even after integrating HiFi and non-HiFi data. We then identified TR size boundaries in the centenarian controls corresponding to the 5th and 95th percentiles of the joint TR allele sizes (2.2 Kbp and 8.4 Kbp, respectively). The number of centenarians with a TR size lower than the 5th percentile was three-fold higher than that of AD cases (1-tailed Fisher’s exact test p=0.023, OR=3.2, *Figure 5E*), and the number of AD cases with a TR size larger than the 95th percentile was two-fold higher than that of centenarians (1-tailed Fisher’s exact test p=0.04, OR=2.0, *Figure 5E*). Given the difficulties in correctly assessing the allele sequences of this TR, we cannot exclude that additional samples suffer from allelic dropouts, especially for the larger expanded allele-sequences.

## 3. Discussion

In this study, we provide novel contributions to better characterise tandem repeats (TRs) with long-read sequencing data. First, we present our novel tools, *otter* and TREAT, that provide a unified workflow to accurately characterise TRs using both Pacific Bioscience (PacBio) and Oxford Nanopore Sequencing Technologies (ONT) datasets. This enabled us to characterise genome-wide TRs in patients with neurodegenerative diseases and genomes from the Human Pangenome Research Consortium (HPRC). Second, we show that in rare instances, long-read sequencing technologies can suffer from abnormal coverage drops in TRs due to potential systematic challenges, particularly in PacBio’s HiFi technology. These coverage drops can lead to TR mis-genotyping, as we observed in CANVAS and Alzheimer’s disease (AD)-associated TRs. Finally, we applied TREAT/*otter* to a case-control setting and replicated TRs previously associated with AD across 494 long-read sequenced AD patients and cognitively healthy centenarian genomes.

Our benchmark of *otter* and TREAT highlighted state-of-the-art performances of our tools in terms of TR genotyping and motif identification accuracy. We showed that *otter,* TREAT, and other existing tools provide generally accurate characterisations of TRs on both PacBio and ONT datasets, and with improved accuracies at higher sequencing coverages. Across technologies, our benchmark revealed that PacBio leads to generally more accurate genotypes for relatively smaller TRs, with PacBio and ONT having similar performances for TRs ranging 500-1000 bp, and ONT leading to more accurate genotypes for larger TRs. These results remained when using other distance metrics as well as in a similar benchmark using the CHM13 reference genome and a larger set of genome-wide TRs.

Our systematic analysis of coverage drops revealed that overall, coverage drops of TRs are rare (0.6%), and do not impact the overall genotyping performances of TREAT/*otter* and other tools. However, our analysis relied on HG002, a highly homozygous genome sequenced at high coverage (38x). Hence, TR coverage drops may be more prevalent in other (low-coverage) genomes that harbour expanded TR sequences, especially those with GC-rich sequences. TRs with coverage drops were often large (>500 bp), high in GC-content, and with higher densities of predicted G-quadruplex DNA secondary structures (G4s). G4s have been previously reported to reduce polymerase efficiency.^40^ As PacBio’s HiFi technology relies on multiple successful passes of a DNA polymerase in a circular DNA template,^20^ we speculate that the interference of G4s might reduce the number of passes in the circular template, possibly leading to lower quality reads (non-HiFi reads). Altogether, incidents of TR coverage drops were enriched in PacBio’s Revio and Sequel 2 datasets, and to a lower extent in ONT’s Duplex and Simplex datasets, with ONT Simplex suffering the least. Although rare, we showed and experimentally validated that coverage drops in TRs can occur at clinically relevant TRs, requiring extra attention when characterising these TRs. To this end, we showed that local vs. global coverage ratio is an effective way to identify such problematic regions, and that for PacBio, these regions can be (in part) rescued by adding noisier non-HiFi data, as shown for the TRs in *ABCA7* and *RFC1* genes.

TREAT and *otter* can be used to genotype and characterise potentially any type of repetitive sequences. However, this remains challenging for very large TRs spanning several kilobases, for example those in telomeric and centromeric regions of the genome. We also note that regions where sequencing error-rates exceed inter-allele dissimilarities may still be difficult to genotype. As the error rate in ONT Simplex data is relatively higher than PacBio and ONT Duplex, this is likely driving the lower genotyping accuracy observed in ONT Simplex. These limitations are not only specific to TREAT and *otter*, but extend to other existing tools. With newer sequencing technologies bringing longer read lengths (*e.g.*, ONT ultra-long reads), together with more complete reference genome assemblies, it might become possible to genotype any satellite region (micro-, mini-, and macro-satellites) in the genome with TREAT and *otter*.

We were able to replicate previously reported TRs associated with AD by comparing a cohort of AD patients and cognitively healthy centenarians. We acknowledge that these TRs were previously identified using different experimental methods (*e.g.* short-read sequencing, southern blotting), and analyses strategies (logistic regressions, linear mixed models, fisher’s exact test).^12,38,39^ While this heterogeneity hampers the direct comparison of the effect size estimates, all associations we observed were in the same direction as the original studies. In particular, the TR intronic of *ABCA7* was shown to carry an odds ratio for AD of 4.5 when one allele was expanded >5.7 Kbp.^12^ Similarly, we observed that individuals carrying larger allele-sequences were significantly associated with AD. However, in our cohort, the effect was mainly driven by cognitively healthy centenarians having a shorter joint-allele size (*i.e.* more AD-protection), rather than AD cases having a more expanded TR-sizes. While we cannot exclude that we have missed some expanded genotypes due to allele dropouts, the centenarians that we included were previously shown to be enriched with the protective alleles in the majority of Single Nucleotide Polymorphisms (SNPs) associated with AD.^41^

In summary, *otter* and TREAT are flexible and accurate bioinformatics tools compatible with different sequencing platforms and requiring minimal input requirements, that enable end-to-end analysis and comparisons of tandem repeats in human genomes with broad applications in research and clinical fields.

## 4. Methods

### 4.1 TREAT

The main analysis is the *assembly* analysis, which uses *otter* for TR genotyping, and is followed by TR content characterisation (identification of motif and number of copies) on the individual TR alleles. In addition to the *assembly* analysis, TREAT implements a *reads* analysis. Here, TR genotyping is performed using an iterative clustering framework based on TR sizes (*Supplementary Methods*). This is followed by TR content characterisation, which is done on all individual reads (*Supplementary Methods* and *Supplementary Results*). This analysis may be preferred when information from all reads is needed, for example for performing a multiple sequence alignment, or when studying somatic instability.

In all cases, TR content characterisation is performed with *pytrf* (https://github.com/lmdu/pytrf). When multiple motif annotations for the same sequence are found by *pytrf*, a consensus representation of the repeat content is generated. Briefly, if the fraction of sequence annotated with a given motif is >95%, then the relative motif is regarded as the best motif describing the TR. In case two or more motifs are found, each describing a portion of the sequence, then the intersection is calculated by intersecting the motif-specific start and end positions. If the intersection is <90%, then the motifs and the relative number of copies are combined. For example, for sequence TGTGTGTGTGTGTGGAGAGAGAGAGAGA, *pytrf* identifies (*i*) 7 copies of TG (ranging positions 1-14, 50% of the sequence covered), and (*ii*) 7 copies of GA (ranging positions 15-28, 50% of the sequence covered). In this case, the combined sequence annotation will be TG+GA, repeated 7+7 times (see *Supplementary Methods*).

TREAT’s analysis module consists of an outlier-detection framework, and a case-control analysis. The outlier-detection scores extreme variations in TR allele sizes across a set of samples. Outliers are detected using a normalised distance that quantifies how far each allele size is from the median allele size, scaled by the variability of the data (*Supplementary Methods*). A p-value for each individual is then calculated by comparing each data point’s distance to a chi-squared distribution. The case-control analysis employs logistic regression models to compare allele sizes (short allele, long allele, and joint allele size) between cases and controls.

### 4.2 *Otter:* a stand-alone, fast, local assembler

*Otter* is a generic stand-alone method for generating fast local assemblies of a given region or genotyping whole-genome *de novo* assemblies. *Otter* in the main genotyping engine of TREAT *assembly* analysis. Briefly, given a region of interest, *otter* uses the htslib library to identify spanning reads (region of interest is fully contained in the reads) and non-spanning reads (only partially contained) in a given BAM file, and extracts the corresponding subsequence per read based on their alignment (*Figure 1B*).^42^ When a reference genome is provided, it will perform local read-realignments on non-spanning reads if it detects a clipping-signal, which can indicate suboptimal mappings to due highly divergent sequences (*Figure 1B*). This is done by aligning (using WFA2-lib alignment library)^43^ the flanking sequences of a region (100 bp by default, modifiable with ‘--flank-size’ parameter) derived from the reference genome onto each read, which are then used to recalibrate the corresponding subsequence of the region of interest. Recalibrated non-spanning reads are reclassified as *spanning* if both flanking sequences are successfully aligned with a minimum length and sequence similarity (by default, 90% sequence similarity, modifiable with ‘--min-sim’ parameter). In the context of TRs, this realignment procedure often correctly recalibrates the alignments of TRs with major length and/or motif-composition differences relative to a reference genome.

*Otter* identifies unique allele-sequences by clustering spanning-reads via pairwise-sequence alignment (*Figure 1B* and *Supplementary Methods*). To manage high somatic variation and/or sequencing errors, *otter* estimates local baseline error-rates per region using a gaussian-kernel density estimator. This produces a one-dimensional distribution of spanning pairwise-sequence distances. In single homozygous allele-sequences, the distribution is unimodal centred at 0. With multiple allele-sequences, the distribution is multimodal, where peaks represent sequence errors between reads from different allele-sequences. *Otter* identifies these peaks and performs hierarchical clustering, stopping when distances exceed the densest peak, partitioning reads into initial clusters. This procedure is followed by a curation step to ensure sufficient read support, adapting to local coverage (*Figure 1B*). If no maximum number of alleles (*α*) is enforced, *otter* outputs all clusters. Otherwise, clusters below the coverage threshold are merged, and if clusters exceed *α*, hierarchical clustering continues until *α* clusters remain. *Otter* then generates a final consensus sequence per cluster via pseudo-partial order alignment procedure of spanning and non-spanning reads inspired from *Ye and Ma, 2016*.^44^

### 4.3 Genomes included for testing

#### HPRC

Publicly available PacBio long-read HiFi data of 47 individuals from the Human Pangenome Reference Consortium (HPRC) were downloaded (*Data Accession*).^34^ For the well characterised HG002 genome,^32^ we also downloaded data generated with Oxford Nanopore (ONT, Duplex and Simplex chemistries) and PacBio Revio technologies. Finally, we generated long-read sequencing data for HG002 using the PacBio Sequel 2 instrument across three SMRT cells, keeping both HiFi and non-HiFi data. ONT data was aligned to the reference genomes (GRCh38 and CHM13) using minimap2 (2.21-r1071, specifying -x map-ont).^45^ PacBio data was aligned using pbmm2 (1.9.0, specifying –preset CCS and –preset SUBREADS respectively for HiFi and non-HiFi data).^20^

#### 100-plus Study cohort and Alzheimer Dementia Cohort

For the replication of TRs previously associated with Alzheimer’s Disease (AD), we used HiFi sequencing (Sequel 2) data from the blood DNA of N=246 patients with AD from the Amsterdam Dementia Cohort (ADC),^36,46^ and N=248 cognitively healthy centenarians from the 100-plus Study cohort.^36,47^ Ten cognitively healthy centenarians were sequenced as a trio, including the blood-derived DNA from the centenarian, the brain-derived DNA from the centenarian and blood-derived DNA from a child of the centenarian. The combined set of a centenarian and child is referred to as parent-child duo throughout the manuscript. Sequencing data pre-processing was conducted as previously described (*Supplementary Methods*).^36^

#### CANVAS patients

We used the HiFi data (Sequel 2) of two patients diagnosed with CANVAS (Cerebellar ataxia with neuropathy and vestibular areflexia syndrome), caused by a TR expansion in *RFC1* gene.^35^

### 4.4 Evaluating *otter* and TREAT performances

#### Comparison with existing tools

We compared TREAT/*otter* to TRGT and LongTR.^22,31^ For the comparison, we used the HG002 genome and a set of 161,382 TRs from PacBio’s repeat catalogue (version 0.3.0, available at https://github.com/PacificBiosciences/trgt/tree/main/repeats). We compared the tools’ genotyped alleles to the expected alleles from the T2T assembly of HG002. As metrics, we used (*i*) normalised edit distance, (*ii*) raw edit distance, (*iii*) allele size correlation between the observed and expected alleles, and (*iv*) fraction of perfectly genotyped alleles. In addition, we evaluated motif identification accuracy, and computational resources.

#### TREAT/otter applications

We compared the performances of TREAT *assembly* and *reads* analyses by correlating the estimated TR allele sizes with each other (*Supplementary Results*). Then, we used TRs for a population stratification analysis: using the set of 161K TRs, we selected the top 20% most variable TRs based on the coefficient of variation (ratio of standard deviation to the mean TR joint allele size). Then we applied Principal Component Analysis (PCA) based on the joint allele sizes. For 40/47 matching samples with Single Nucleotide Polymorphisms (SNP) data from the 1000Genome project,^48^ we also performed PCA based on 30,544 randomly sampled common (minor allele frequency >10%) SNPs.

To evaluate clinical applicability, we applied the TREAT/*otter* outlier analysis module on the combined dataset of 47 HPRC genomes plus the two CANVAS patients and the ten parent-child duos. For this analysis, we focused on 35 clinically relevant TRs (*Table S1*), that were previously associated with neurological diseases.^7,8,12^ Finally, TREAT/*otter* case-control analysis module was used to replicate the association of four TRs that were previously associated with Alzheimer’s Disease (AD).^12,38,39^ The commands used for the outlier and case-control analyses are available in *Supplementary Methods*.

### 4.5 Systematic analysis of allele dropouts in tandem repeats

#### Curated set of TRs in CHM13

We downloaded and curated repeat annotations for the CHM13 reference genome (version 2.0, https://github.com/marbl/CHM13, *Supplementary Methods*). This curated dataset counted 864,424 TRs genome-wide. We extracted the corresponding parental and maternal allele-sequences in HG002 for these TRs by aligning the HG002 T2T assembly (version 0.7) to CHM13.^32^

#### TRs unique to CHM13

We first genotyped the 864K TRs using *otter* in HG002 from different technologies (Sequel 2, Revio, Simplex and Duplex), and at different coverage levels (5x, 10x, 15x, 20x, 25x and 30x), and calculated the normalised edit distance between observed and expected TR alleles (*Supplementary Results*). We then focussed on a set of TRs present in CHM13 and absent in GRCh38, and used TREAT/*otter* to characterise the repeat content of these TRs in 47 genomes from HPRC.

#### Evaluation of coverage drops in TR

Using HG002 data from Sequel 2, Revio, Simplex and Duplex technologies (∼30x coverage each), we calculated the ratio between local TR coverage and average global coverage. TRs where this ratio was <0.25 were regarded as low-coverage TRs. We then investigated sequence characteristics of low-coverage TR, including average size, dinucleotide content, and propensity to form G-quadruplex DNA secondary structures (G4s). For the latter, we used pqsfinder (v2.10.1) with ‘min_score = 20’ parameter.^49^

## Data access

Human Pangenome Consortium data is publicly available and can be downloaded from https://github.com/human-pangenomics/HPP_Year1_Data_Freeze_v1.0?tab=readme-ov-file.

Long-read sequencing data generated with PacBio Sequel 2 for the 2 CANVAS patients as well as 246 AD patients and 248 cognitively healthy centenarians is available upon submission of a research proposal to the Alzheimer Genetics Hub (AGH, https://alzheimergenetics.org/).

## Consent statement

The Medical Ethics Committee of the Amsterdam UMC and Radboud UMC approved all studies. All participants and/or their legal representatives provided written informed consent for participation in clinical and genetic studies.

## Funding

Part of the work in this manuscript was carried out on the Cartesius supercomputer, which is embedded in the Dutch national e-infrastructure with the support of SURF Cooperative. Computing hours were granted to H. H. by the Dutch Research Council (100plus: project# vuh15226, 15318, 17232, and 2020.030; Role of VNTRs in AD; project# 2022.31, Alzheimers Genetics Hub project# 2022.38). N.T is appointed at ABOARD and H.H, M.R, S.L are recipients of ABOARD, a public-private partnership receiving funding from ZonMW (#73305095007) and Health∼Holland, Topsector Life Sciences & Health (PPP-allowance; #LSHM20106).^41,50^ This work is supported by a VIDI grant from the Dutch Scientific Counsel (#NWO 09150172010083) and a public-private partnership with TU Delft and PacBio, receiving funding from ZonMW and Health∼Holland, Topsector Life Sciences & Health (PPP-allowance), and by Alzheimer Nederland WE.03-2018-07. S.L. is recipient of ZonMW funding (#733050512). H.H. was supported by the Hans und Ilse Breuer Stiftung (2020), Dioraphte 16020404 (2014) and the HorstingStuit Foundation (2018). Acquisition of the PacBio Sequel 2 long read sequencing machine was supported by the ADORE Foundation (2022).

## Conflicts of interest

All authors declare no conflict of interest.

## Acknowledgements

We are grateful to all the reviewers involved in the peer-review process for their comments which have largely improved our manuscript.

## Notes

### Competing Interest Statement

The authors have declared no competing interest.

### Summary of Updates

This version of the manuscript has been revised to update the manuscript with the latest analyses that were requested during the peer-review process. These analysis include a formal benchmarking comparison or the newly developed tools with current state-of-the-art tools in long-read sequencing data.

## References

1. Hannan, A. J. Tandem repeats mediating genetic plasticity in health and disease. Nat Rev Genet 19, 286–298 (2018).

2. Lynch, M. et al. A genome-wide view of the spectrum of spontaneous mutations in yeast. Proc Natl Acad Sci U S A 105, 9272–9277 (2008).

3. Pearson, C. E., Edamura, K. N. & Cleary, J. D. Repeat instability: mechanisms of dynamic mutations. Nat Rev Genet 6, 729–742 (2005).

4. Subramanian, S., Mishra, R. K. & Singh, L. Genome-wide analysis of microsatellite repeats in humans: their abundance and density in specific genomic regions. Genome Biol 4, R13 (2003).

5. Linthorst, J. et al. Extreme enrichment of VNTR-associated polymorphicity in human subtelomeres: genes with most VNTRs are predominantly expressed in the brain. Transl Psychiatry 10, 369 (2020).

6. Dumbovic, G., Forcales, S.-V. & Perucho, M. Emerging roles of macrosatellite repeats in genome organization and disease development. Epigenetics 12, 515–526 (2017).

7. McMurray, C. T. Mechanisms of trinucleotide repeat instability during human development. Nat Rev Genet 11, 786–799 (2010).

8. Khristich, A. N. & Mirkin, S. M. On the wrong DNA track: Molecular mechanisms of repeat-mediated genome instability. J Biol Chem 295, 4134–4170 (2020).

9. Stevanovski, I. et al. Comprehensive genetic diagnosis of tandem repeat expansion disorders with programmable targeted nanopore sequencing. Sci Adv 8, eabm5386 (2022).

10. Yu, S. et al. Fragile X Genotype Characterized by an Unstable Region of DNA. Science 252, 1179–1181 (1991).

11. DeJesus-Hernandez, M. et al. Expanded GGGGCC hexanucleotide repeat in noncoding region of C9ORF72 causes chromosome 9p-linked FTD and ALS. Neuron 72, 245–256 (2011).

12. On Behalf of the BELNEU Consortium et al. An intronic VNTR affects splicing of ABCA7 and increases risk of Alzheimer’s disease. Acta Neuropathologica 135, 827–837 (2018).

13. De Roeck, A., Van Broeckhoven, C. & Sleegers, K. The role of ABCA7 in Alzheimer’s disease: evidence from genomics, transcriptomics and methylomics. Acta Neuropathol 138, 201–220 (2019).

14. Dolzhenko, E. et al. ExpansionHunter: a sequence-graph-based tool to analyze variation in short tandem repeat regions. Bioinformatics 35, 4754–4756 (2019).

15. Gymrek, M., Golan, D., Rosset, S. & Erlich, Y. lobSTR: A short tandem repeat profiler for personal genomes. Genome Res. 22, 1154–1162 (2012).

16. Kristmundsdóttir, S., Sigurpálsdóttir, B. D., Kehr, B. & Halldórsson, B. V. popSTR: population-scale detection of STR variants. Bioinformatics 33, 4041–4048 (2017).

17. Bakhtiari, M., Shleizer-Burko, S., Gymrek, M., Bansal, V. & Bafna, V. Targeted genotyping of variable number tandem repeats with adVNTR. Genome Res. 28, 1709–1719 (2018).

18. Gelfand, Y., Hernandez, Y., Loving, J. & Benson, G. VNTRseek-a computational tool to detect tandem repeat variants in high-throughput sequencing data. Nucleic Acids Res 42, 8884–8894 (2014).

19. Eslami Rasekh, M., Hernández, Y., Drinan, S. D., Fuxman Bass, J. I. & Benson, G. Genome-wide characterization of human minisatellite VNTRs: population-specific alleles and gene expression differences. Nucleic Acids Research 49, 4308–4324 (2021).

20. Wenger, A. M. et al. Accurate circular consensus long-read sequencing improves variant detection and assembly of a human genome. Nat Biotechnol 37, 1155–1162 (2019).

21. Sereika, M. et al. Oxford Nanopore R10.4 long-read sequencing enables the generation of near-finished bacterial genomes from pure cultures and metagenomes without short-read or reference polishing. Nat Methods 19, 823–826 (2022).

22. Dolzhenko, E. et al. Characterization and visualization of tandem repeats at genome scale. Nat Biotechnol (2024) doi:10.1038/s41587-023-02057-3.

23. Chiu, R., Rajan-Babu, I.-S., Friedman, J. M. & Birol, I. Straglr: discovering and genotyping tandem repeat expansions using whole genome long-read sequences. Genome Biol 22, 224 (2021).

24. Masutani, B., Kawahara, R. & Morishita, S. Decomposing mosaic tandem repeats accurately from long reads. Bioinformatics 39, btad185 (2023).

25. Gustafson, J. A. et al. Nanopore Sequencing of 1000 Genomes Project Samples to Build a Comprehensive Catalog of Human Genetic Variation. http://medrxiv.org/lookup/doi/10.1101/2024.03.05.24303792 (2024) doi:10.1101/2024.03.05.24303792.

26. Ren, J., Gu, B. & Chaisson, M. J. P. vamos: variable-number tandem repeats annotation using efficient motif sets. Genome Biol 24, 175 (2023).

27. Gouil, Q. & Keniry, A. Latest techniques to study DNA methylation. Essays in Biochemistry 63, 639–648 (2019).

28. Liu, Y. et al. DNA methylation-calling tools for Oxford Nanopore sequencing: a survey and human epigenome-wide evaluation. Genome Biol 22, 295 (2021).

29. Amarasinghe, S. L. et al. Opportunities and challenges in long-read sequencing data analysis. Genome Biol 21, 30 (2020).

30. Mirkin, S. M. Expandable DNA repeats and human disease. Nature 447, 932–940 (2007).

31. Ziaei Jam, H., et al. LongTR: genome-wide profiling of genetic variation at tandem repeats from long reads. Genome Biol 25, 176 (2024).

32. Jarvis, E. D. et al. Semi-automated assembly of high-quality diploid human reference genomes. Nature 611, 519–531 (2022).

33. Baid, G. et al. DeepConsensus improves the accuracy of sequences with a gap-aware sequence transformer. Nat Biotechnol (2022) doi:10.1038/s41587-022-01435-7.

34. Wang, T. et al. The Human Pangenome Project: a global resource to map genomic diversity. Nature 604, 437–446 (2022).

35. van de Pol, M. et al. Detection of the ACAGG Repeat Motif in RFC1 in Two Dutch Ataxia Families. Mov Disord 38, 1555–1556 (2023).

36. Salazar, A., et al. An AluYb8 Retrotransposon Characterises a Risk Haplotype of TMEM106B Associated in Neurodegeneration. http://medrxiv.org/lookup/doi/10.1101/2023.07.16.23292721 (2023) doi:10.1101/2023.07.16.23292721.

37. Zhang, Y., et al. MotifScope: A Multi-Sample Motif Discovery and Visualization Tool for Tandem Repeats. http://biorxiv.org/lookup/doi/10.1101/2024.03.06.583591 (2024) doi:10.1101/2024.03.06.583591.

38. Guo, M. H., Lee, W.-P., Vardarajan, B., Schellenberg, G. D. & Phillips-Cremins, J. Polygenic burden of short tandem repeat expansions promote risk for Alzheimer’s disease. medRxiv 2023.11.16.23298623 (2023) doi:10.1101/2023.11.16.23298623.

39. Wang, H. et al. Structural Variation Detection and Association Analysis of Whole-Genome-Sequence Data from 16,905 Alzheimer’s Diseases Sequencing Project Subjects. medRxiv 2023.09.13.23295505 (2023) doi:10.1101/2023.09.13.23295505.

40. Lago, S. et al. Promoter G-quadruplexes and transcription factors cooperate to shape the cell type-specific transcriptome. Nat Commun 12, 3885 (2021).

41. Tesi, N. et al. Cognitively healthy centenarians are genetically protected against Alzheimer’s disease. Alzheimer’s & Dementia 20, 3864–3875 (2024).

42. Bonfield, J. K. et al. HTSlib: C library for reading/writing high-throughput sequencing data. Gigascience 10, giab007 (2021).

43. Marco-Sola, S. et al. Optimal gap-affine alignment in *O* (*s*) space. Bioinformatics 39, btad074 (2023).

44. Ye, C. & Ma, Z. (Sam). Sparc: a sparsity-based consensus algorithm for long erroneous sequencing reads. PeerJ 4, e2016 (2016).

45. Li, H. Minimap2: pairwise alignment for nucleotide sequences. Bioinformatics 34, 3094– 3100 (2018).

46. van der Flier, W. M. & Scheltens, P. Amsterdam Dementia Cohort: Performing Research to Optimize Care. Journal of Alzheimer’s Disease 62, 1091–1111 (2018).

47. Holstege, H. et al. The 100-plus Study of cognitively healthy centenarians: rationale, design and cohort description. European Journal of Epidemiology (2018) doi:10.1007/s10654-018-0451-3.

48. 1000 Genomes Project Consortium et al. A global reference for human genetic variation. Nature 526, 68–74 (2015).

49. Hon, J., Martínek, T., Zendulka, J. & Lexa, M. pqsfinder: an exhaustive and imperfection-tolerant search tool for potential quadruplex-forming sequences in R. Bioinformatics 33, 3373–3379 (2017).

50. Dreves, M. A. E. et al. Rationale and design of the ABOARD project (A Personalized Medicine Approach for Alzheimer’s Disease). Alzheimers Dement (N Y) 9, e12401 (2023).

