## Supplementary Text, Figures and Tables for "Characterising tandem repeat complexities across long-read sequencing platforms with TREAT and *otter*": Supplements - Clean PDF.pdf

### Table of Contents

**1. Supplementary Results** **2**

1.1 Comparison of *otter* and TREAT with existing tools 2

1.2 TREAT's unified workflow enables diverse characterisations of tandem repeats 3

1.3 Tandem repeats may be sensitive to coverage dropouts in long-read sequencing 4

**2. Supplementary Figures** **7**

**3. Supplementary Methods** **19**

3.1 Long-read sequencing with PacBio Sequel II and publicly available data from ONT and PacBio instruments 19

3.2 Comparison of *otter* and TREAT with existing tools 19

3.3 TREAT 21

3.4 *Otter* 23

3.5 Cohort and sequencing details 24

3.6 Southern Blotting 25

**4. Supplementary Tables** **27**

Table S1: TR associated with neurological diseases 27

Table S2: TR previously associated with AD 29

**5. References** 30

### 1. Supplementary Results

#### 1.1 Comparison of *otter* and TREAT with existing tools

*Normalised edit distance between PacBio Sequel 2 and Revio*: In general, the normalised edit distance was lower for PacBio's Sequel 2 data in comparison with Revio data: error rates ranged 0.1-2.3% (*otter*) and 0.1-3.8% (TRGT) in Sequel 2 data, and ranged 0.2-2.5% (*otter*) and 0.2-3.3% (TRGT) in Revio data (*Figure 2A*). In ONT data, the normalised edit distance was lower for Duplex data in comparison with Simplex data: error rates ranged 0.3-1.8% (*otter*) and 0.2-2.1% (LongTR) in Duplex data, and ranged 0.3-4.9% (*otter*) and 0.3-3.6% (LongTR) in Simplex data. Note that LongTR was not able to genotype ~20% of TRs at coverage 5x (*Figure 2B*).

*Raw edit distance across tools and technologies*: Using raw edit distance as metric, similar results as to using normalised edit distance were obtained. Error rates ranged 0.1-78.3 (*otter*) and 0.2-114.9 (TRGT) in Sequel 2 data, and ranged 0.2-95.7 (*otter*) and 0.3-107.0 (TRGT) in Revio data. In ONT data, error rates ranged 0.4-51.7 (*otter*) and 0.5-96.5 (LongTR) in Duplex data, and ranged 0.6-133.6 (*otter*) and 0.7-105.8 (LongTR) in Simplex data. Note that LongTR was not able to genotype ~20% of TRs at coverage 5x (*Figure S1D-F*).

*Correlation between TR allele sizes across tools and technologies*: When correlating observed and expected TR allele size across tools and technologies, we observed a similar pattern. Average correlation values (Spearman correlation) ranged 0.91-0.99 (*otter*) and 0.90-0.99 (TRGT) for Sequel 2 data; 0.90-0.99 (*otter*) and 0.91-0.99 (TRGT) for Revio data; 0.90-0.99 (*otter*) and 0.90-0.99 (LongTR) for Simplex data; 0.96-0.99 (*otter*) and 0.91-0.99 (LongTR) for Duplex data (*Figure S1A-D*).

### 1.2 TREAT's unified workflow enables diverse characterisations of tandem repeats

*Genotyping accuracy of otter in ~864K TRs from CHM13*: We benchmarked *otter* on HG002 using a larger set of ~864K TRs from CHM13 reference genome. We used all sequencing technologies (PacBio Sequel 2 and Revio, and ONT Simplex and Duplex), and varied sequencing coverages (5x, 10x, 15x, 20x, 25x, and 30x). Across all technologies, increasing coverage yielded more accurate TR-sequence assemblies (*Figure S10*). At 15x coverage, the average sequence dissimilarities were 0.88% and 0.51% for ONT's Duplex and Simplex chemistries, respectively, and 0.40% and 0.29% for PacBios's Revio and Sequel 2, with slight improvements at higher coverages (*Figure S10*). Overall, we found that *otter*'s assemblies were most accurate when using Sequel 2 HiFi data, followed by Revio, Duplex, and Simplex (*Figure S10*). A closer analysis revealed that Sequel 2 HiFi data had significantly lower read-sequence variability relative to Duplex, Simplex, and Revio (Wilcoxon rank-sum test, p-value <2.2e-16). This is possibly due to a combination of increased sequencing errors and somatic variation in Revio data relative to Sequel 2, which ultimately yielded suboptimal assemblies relative to the expected allele-sequences from the HG002 T2T assembly.

*Comparison of TREAT reads and assembly modes*: We compared TREAT *assembly* analysis, which uses *otter* for TR genotyping, to TREAT *reads* analysis, which genotypes TR alleles using a clustering framework based on TR sizes as observed in the individual reads spanning the TR. For this analysis, we used the curated PacBio genome-wide catalogue of 161K TRs, and all 47 genomes from the HPRC. Spearman correlations were calculated for the shorter (in size), longer, and joint alleles (*i.e.* the sum of the shorter and longer alleles). We found nearly perfect concordance between the *assembly* and *reads* analyses when correlating the size of the TRs across the shorter, longer, and joint allele sizes (Spearman correlation=0.99, p<2.0x10-16 in all cases). This shows that the two analyses deliver correlated results, and their choice depends on the analysis setting: for a large set of TRs, we advise to use *assembly* analysis, while when individual read information is valuable, it is advisable to use the *reads* analysis.

#### 1.3 Tandem repeats may be sensitive to coverage dropouts in long-read sequencing

*Coverage drops in the pathogenic TR in RFC1*: We used TREAT/*otter* to characterise the intronic TR in *RFC1* in 47 genomes from the HPRC consortium, two CANVAS patients, and 10 parent-child duos. We found that CANVAS patient 2 carried a ~7.7 Kbp and a ~8.5 Kbp *RFC1* TR allele-sequence, both largely composed of the disease-associated motif (ACAGG)n, repeated 1489 and 1710 times, respectively. In addition, in the larger allele, the (AACAG)n was also observed (*Figure 3*). CANVAS patient 1 carried a ~6.3 Kbp and a ~7.7 Kbp TR allele-sequence mostly composed of the same (ACAGG) motif, repeated 1270 and 1540 times, respectively (*Figure 3*). Interestingly, although this motif is enriched in non-European individuals, both patients were of Dutch ancestry. This analysis showcases not only TREAT and *otter* utility in characterising tandem-repeats in a clinical setting, but also to facilitate the discovery of novel motif compositions. Although to a lower extent compared to the CANVAS patients, expansions of this TR were also observed in other samples (*Figure 3*). These a heterozygous TR expansion in one parent of the 10 parent-child duos (C7_BLOOD, *Figure 3*). Interestingly, the child of this individual reported homozygous wild-type alleles, which was surprising. The ratio of local *versus* whole-genome coverage for the TR in the parent, using HiFi data alone, was 0.66 (31 total local reads, 47.3x average whole-genome coverage), while for the child it was 0.52 (7 total local reads, 13.5x average whole-genome coverage). Assuming that whole-genome coverage follows a Poisson distribution, these ratios are significantly lower than expected (p-value of 0.00773 and 0.02 for the parent and child, respectively). When we included non-HiFi data, this ratio increased to respectively 1.06 (50 total local reads) and 0.92 (13 total local reads) in the centenarian and child. Furthermore, the expanded allele in the child was supported, and could be rescued (*Figure 4*).

*Coverage drops in the Alzheimer's Disease (AD)-associated TR in ABCA7*: This intronic TR in *ABCA7* was previously identified as a risk factor for Alzheimer's Disease (AD). We further investigated challenges when sequencing this TR by comparing TR-lengths from Southern Blotting assay (*Supplementary Methods*) and long-read sequencing data generated on the Sequel 2 instrument in nine Dutch individuals (coverage ranging 11.5-34x, median read-lengths of 14-19 Kbp). We found similar coverage drops in the *ABCA7* TR in all nine genomes, with local coverages ranging 1-7x. We found low-concordance between lengths in HiFi data and the observed lengths from Southern Blotting assay, suggesting challenges in accurately characterising this TR with HiFi technology. We then used TREAT/*otter* to integrate HiFi and non-HiFi data in the nine centenarian genomes for which experimental validation by Southern Blotting was available. The inclusion of non-HiFi data increased read-support by 4-folds to an average coverage of 22x. Notably, all technologies highlight a potential mis-assembly in HG002 as they do not support the expanded version of the *ABCA7* TR. Manual inspection revealed that the expanded TR in the HG002 assembly harboured TE-like sequences not supported by any of the long-read datasets, suggesting either variation specific to the sequenced HG002 cell-line, or a mis-assembly in the HG002 reference.

*Systematic analysis of coverage drops in TRs*: The average coverage of the 454 low-coverage TRs in PacBio in ONT Simplex was 37.43±16.65, with average local to global coverage ratio of 0.97±0.43. In ONT Duplex, the average coverage was 25.36±12.85, with an average ratio of 0.66±0.33 (≥ 22x coverage, average ratio=0.60). Therefore, the TRs problematic in PacBio instruments were generally well-supported by ONT. In contrast, the 49 problematic regions in ONT reported average coverage of 17.43±9.45 in PacBio Revio, with average ratios of 0.45±0.25, and average coverage of 13.84±13.96 in Sequel 2 data, with an average ratio of

0.38±0.38. 9/20 TRs problematic in ONT had no HIFI reads in both PacBio instruments, and the remaining 11/20 TRs were supported by 6-9 HIFI reads (ratio ranging 0.16-0.24) from one or both instruments, suggesting that these TRs may be challenging to sequence with current long-read sequencing technologies.

### 2. Supplementary Figures

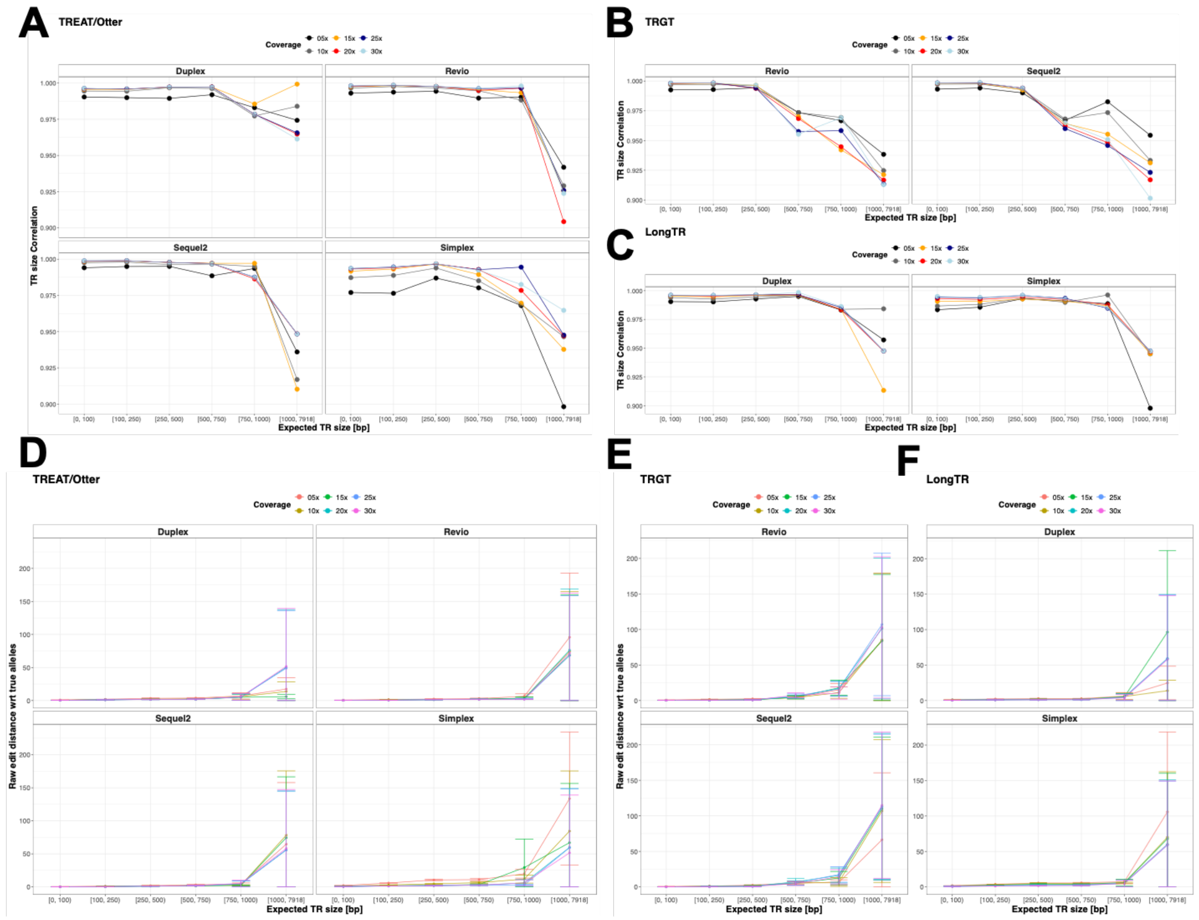


*Figure S1: Additional metrics for TR genotyping accuracy across tools, technologies, and sequencing coverage.* ***A-B-C.*** *Show the correlation between observed and expected allele size, respectively for otter, TRGT, and LongTR.* ***D-E-F.*** *Show the raw edit distance between observed and expected allele sizes, respectively for otter, TRGT, and LongTR.*

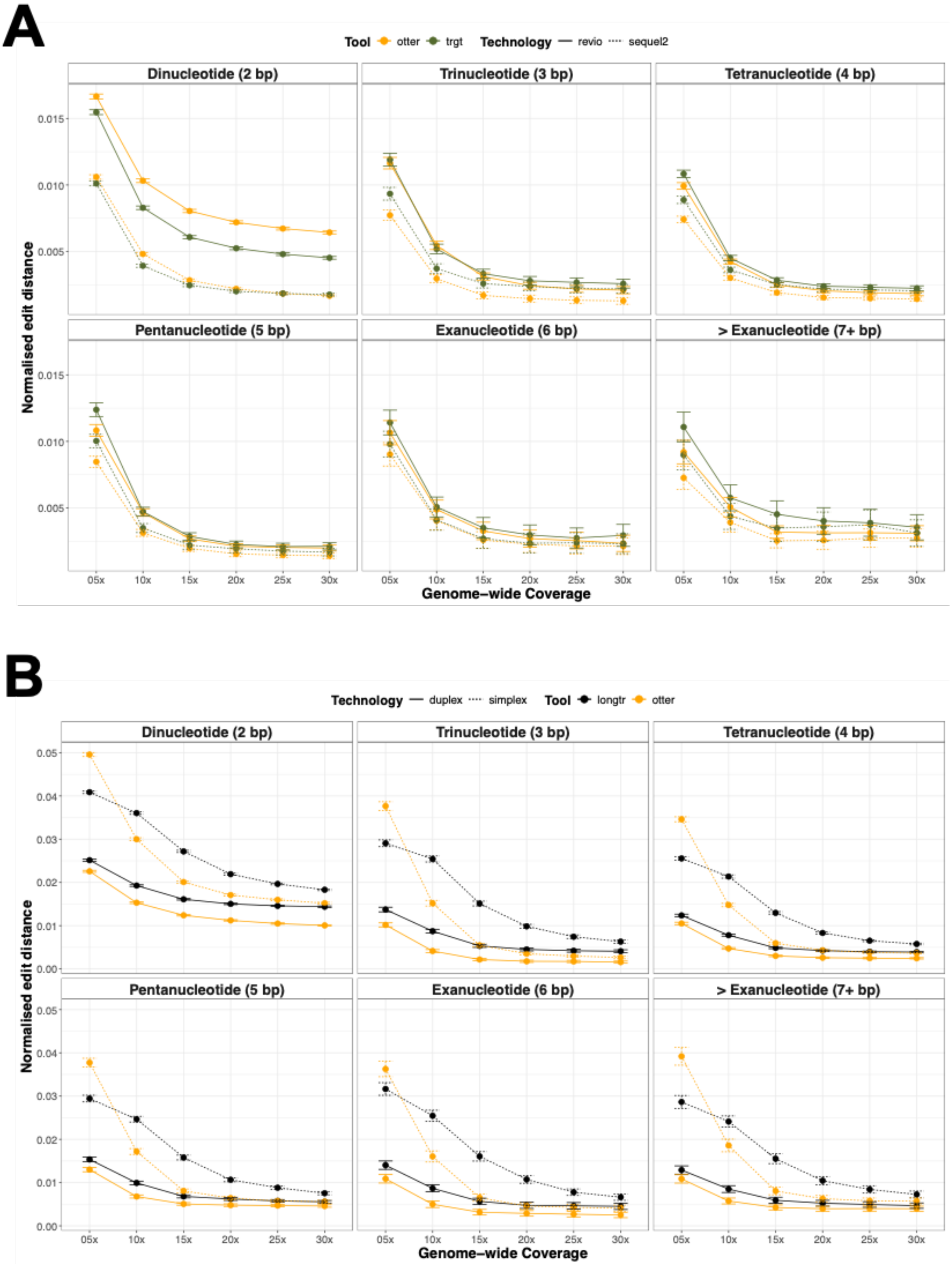


*Figure S2: TR genotyping accuracy in the context of TR motif length.* ***A.*** *Shows the genotyping performances of otter and TRGT in PacBio Sequel 2 and Revio data at different coverage levels and for different TR motif lengths.* ***B.*** *Shows the same performances between otter and*

*LongTR on ONT Simplex and Duplex data. The TR motif size was based on TREAT motif characterization.*

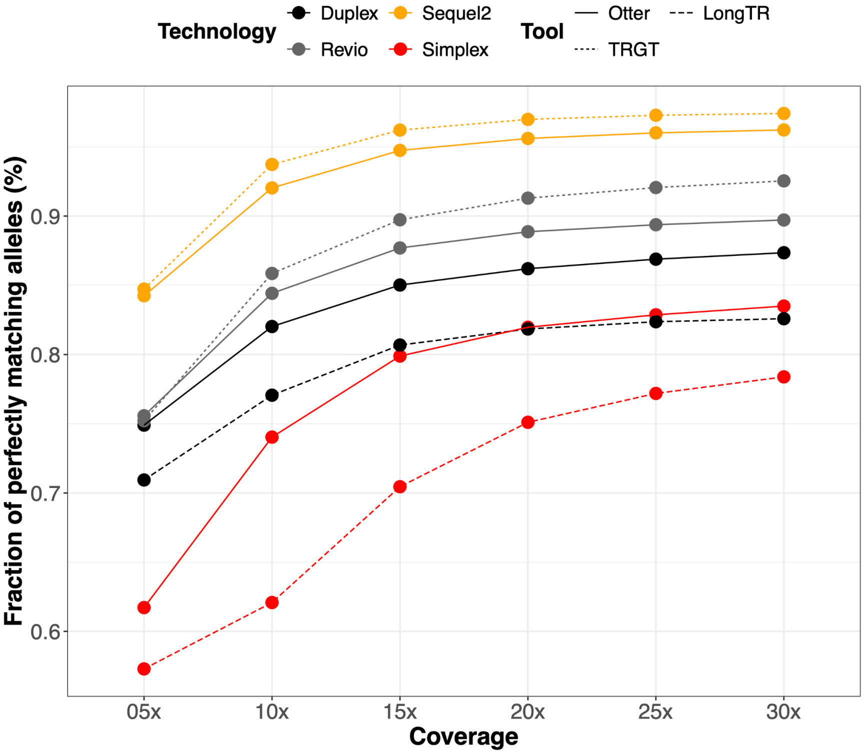


*Figure S3: Fraction of perfectly matched TR alleles across tools, sequencing technologies and coverage levels. For this, we used HG002 data across different coverage levels and sequencing technologies. The Y-axis shows the fraction of alleles with an edit distance of 0 (i.e. perfect matches) between the genotyped and expected alleles.*

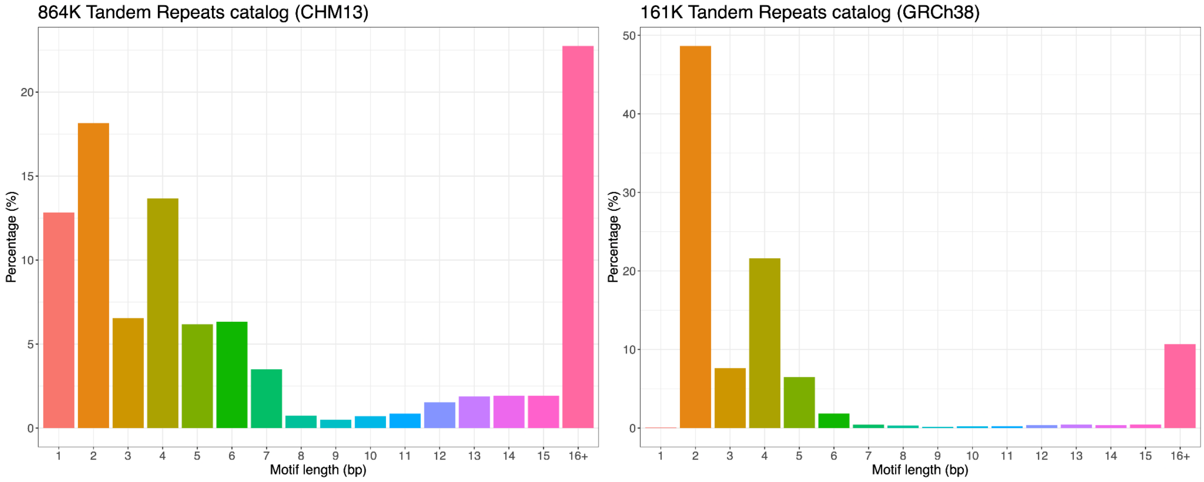

*Figure S4: Motif distribution in the set of 864K repeats in the CHM13 reference genome (left), as well as the 161K tandem repeats in GRCh38 reference genome from PacBio's catalogue of repeats (right).*

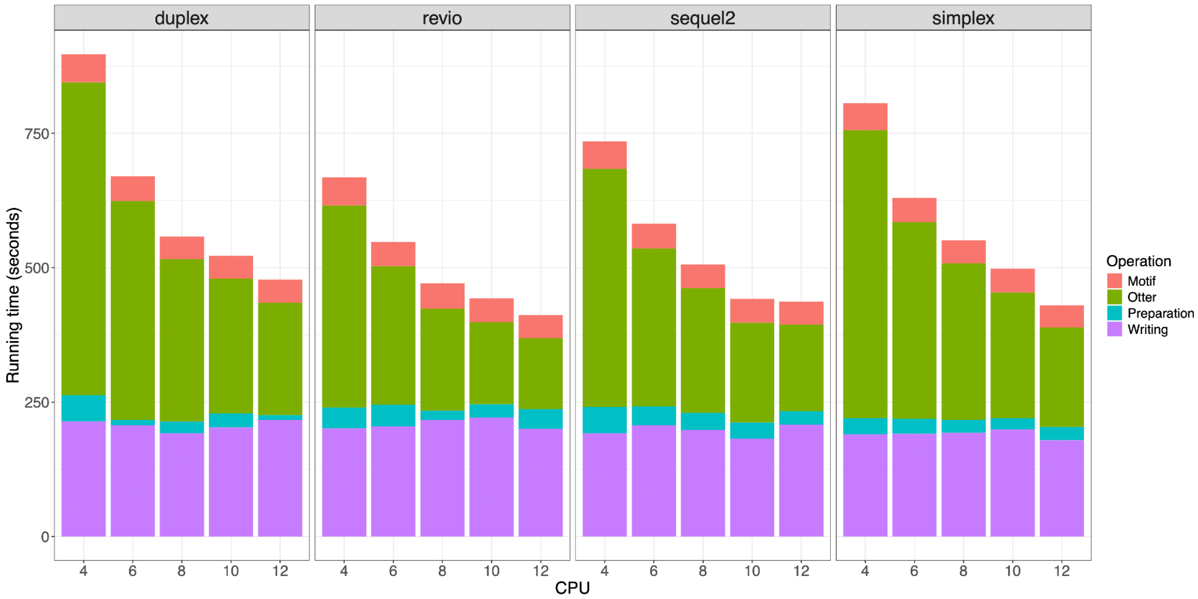


*Figure S5: Multithreading performances of TREAT/otter. X-axis shows the number of CPUs used. Running time is split across internal operations. For this experiment, we used HG002 data (15x coverage) across different sequencing technologies, and 161K TRs from PacBio catalogue.*

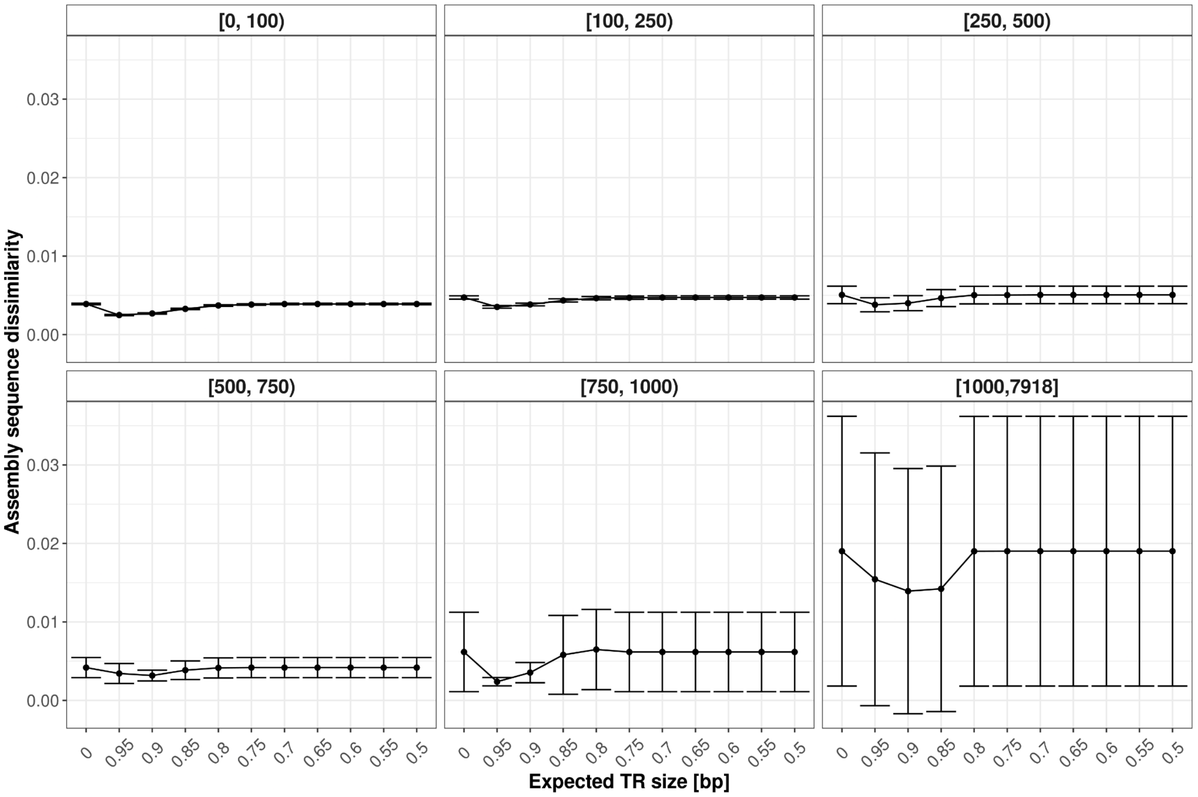


*Figure S6: TR genotyping accuracy when using HiFi + non-HiFi data. This analysis was performed on PacBio Sequel 2 data of HG002 (~38x coverage), for which both HiFi and the complete set of non-HiFi data was available. X-axis shows the read quality of included non-HiFi data (at 0, only HiFi data was included). The different panels refer to TR size.*

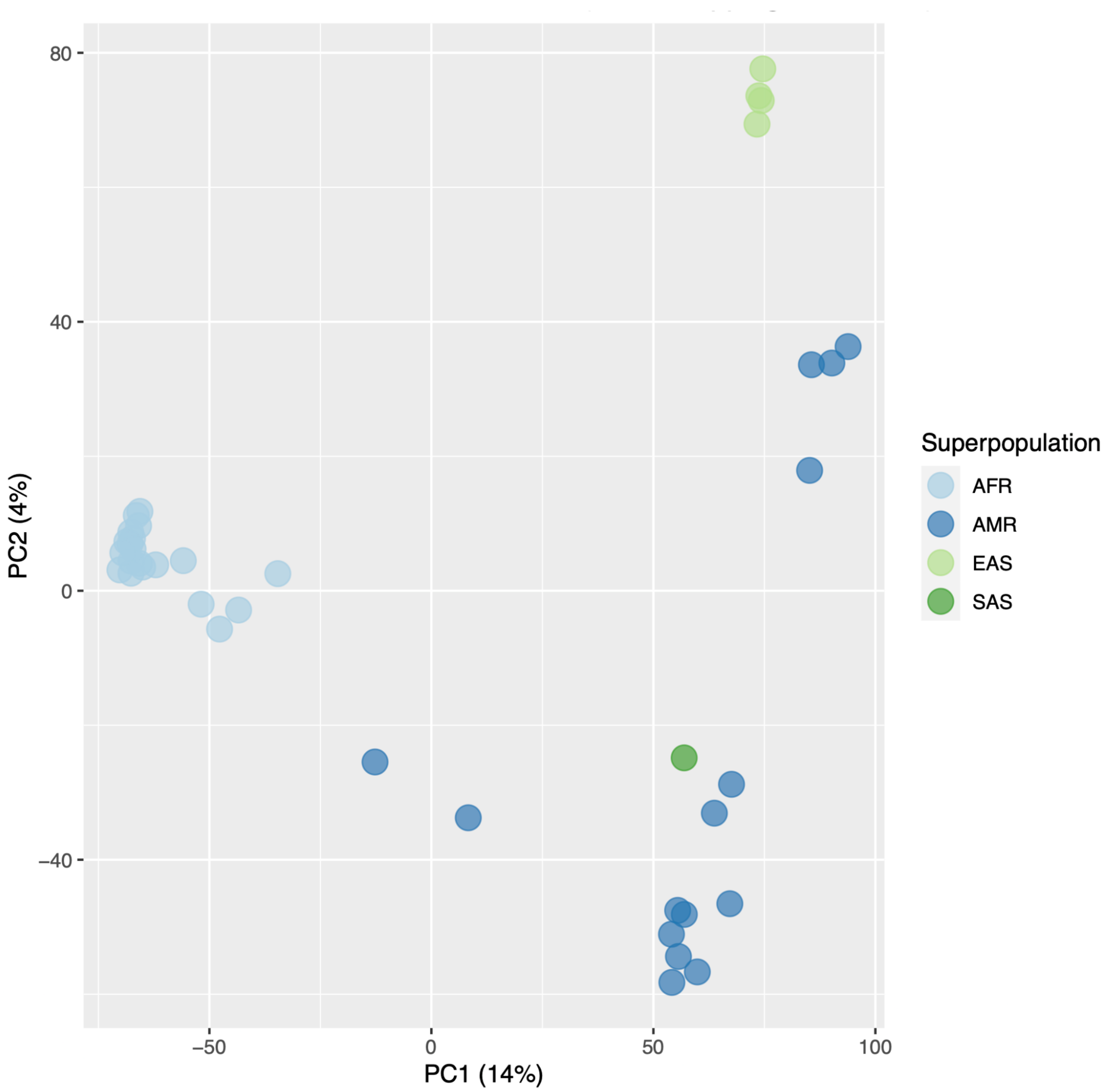


*Figure S7: Population PCA based on single nucleotide polymorphisms (SNPs). We used 40/47 individuals for which both SNP array data and long-read sequencing was available. For the PCA, we used ~30k randomly sampled SNPs with a minor allele frequency >10%.*

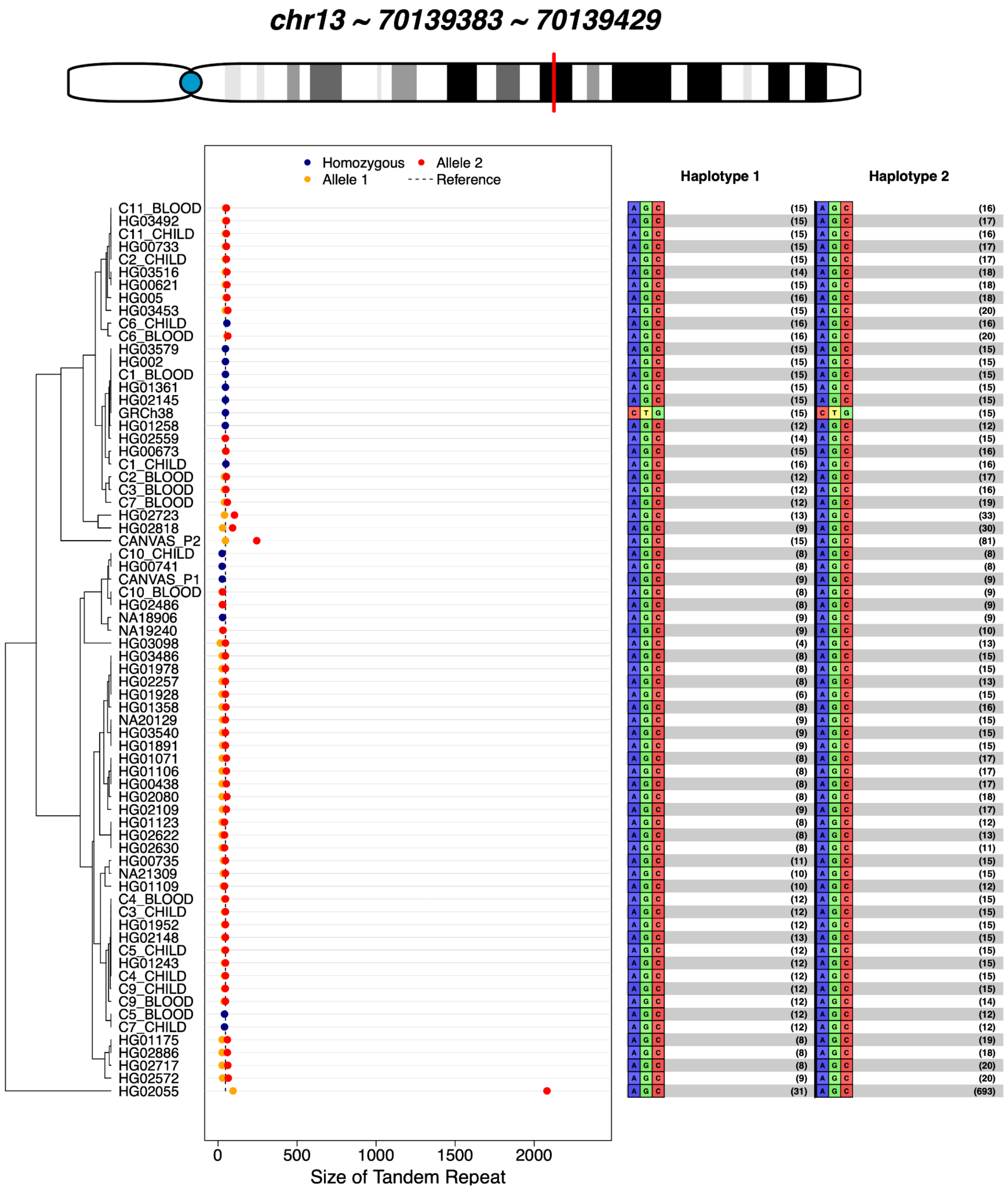


*Figure S8: TREAT plot showing the TR in ATXN8 gene across 47 HPRC samples as well as two CANVAS patients. The left side of the plot shows the TR allele sizes. Red circles identify homozygous genotypes, while Blue/Orange genotypes refer to the shorter and longer alleles (when heterozygous), respectively. Individuals are sorted based on a hierarchical clustering scheme. The right side of the plot shows the motif composition for each of the two alleles, Haplotype 1 and Haplotype 2, along with the relative number of copies.*

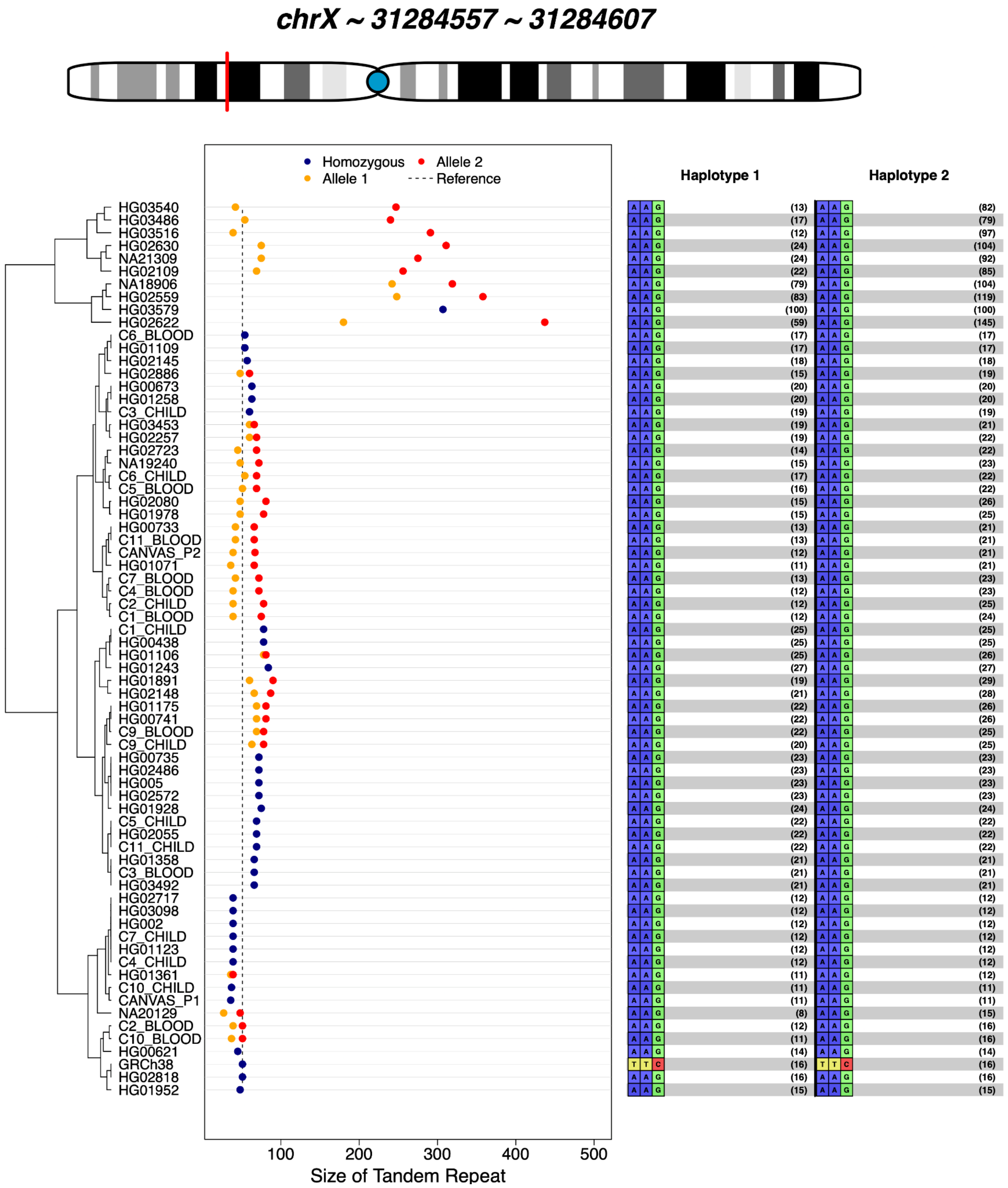


*Figure S9: TREAT plot showing the TR in DMD gene across 47 HPRC samples as well as two CANVAS patients. The left side of the plot shows the TR allele sizes. Red circles identify homozygous genotypes, while Blue/Orange genotypes refer to the shorter and longer alleles (when heterozygous), respectively. Individuals are sorted based on a hierarchical clustering*

*scheme. The right side of the plot shows the motif composition for each of the two alleles, Haplotype 1 and Haplotype 2, along with the relative number of copies.*

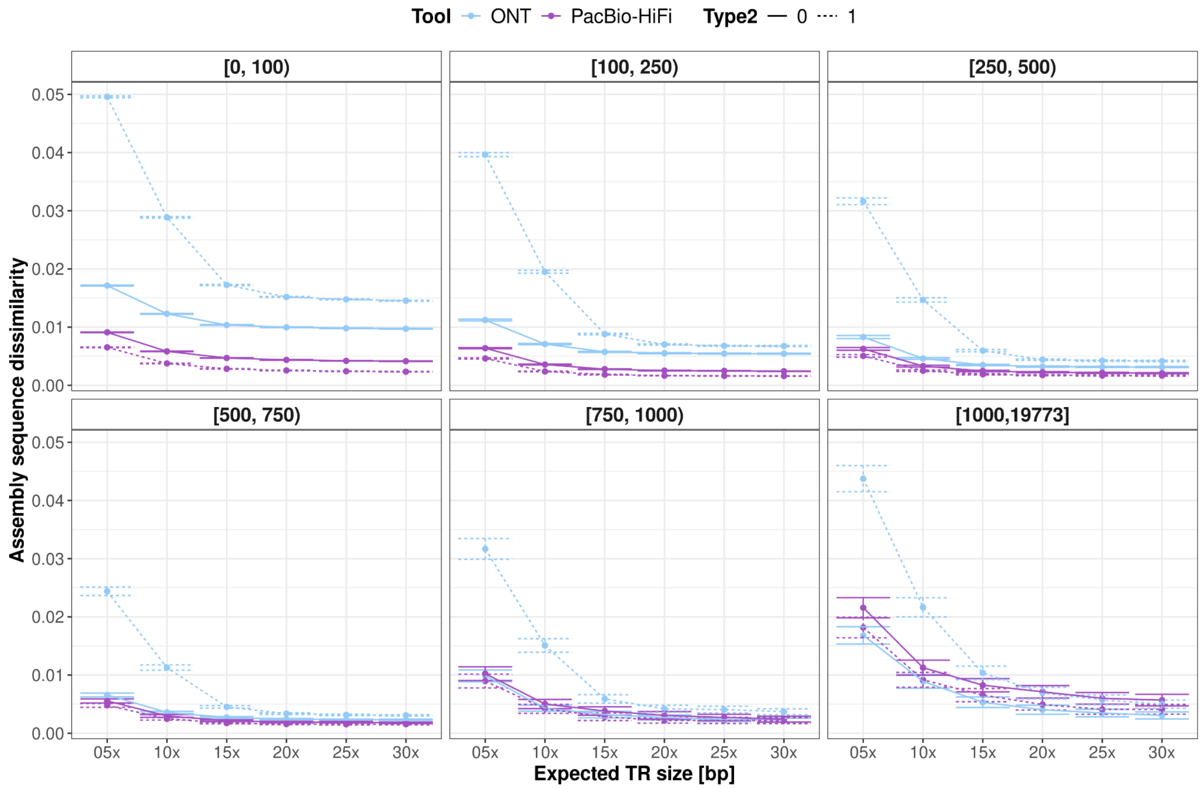


*Figure S10: Otter performances on ~864K TRs from CHM13 reference genome. Solid lines refer to Revio and Simplex, respectively for PacBio and ONT. Dotted lines refer to Sequel 2 and Duplex, respectively for PacBio and ONT.* ***A.*** *Size distribution of the curated set of 864,170 unique TRs (1,727,475 allele-sequences in total), with an average length of 93 bp.* ***B.*** *Assembly-sequence error-rates as a function of the TR-size and sequencing technologies.* ***C.*** *Run time of otter as a function of the coverage and sequencing technologies.* ***D.*** *Memory consumption of otter as a function of the coverage and different sequencing technologies.*

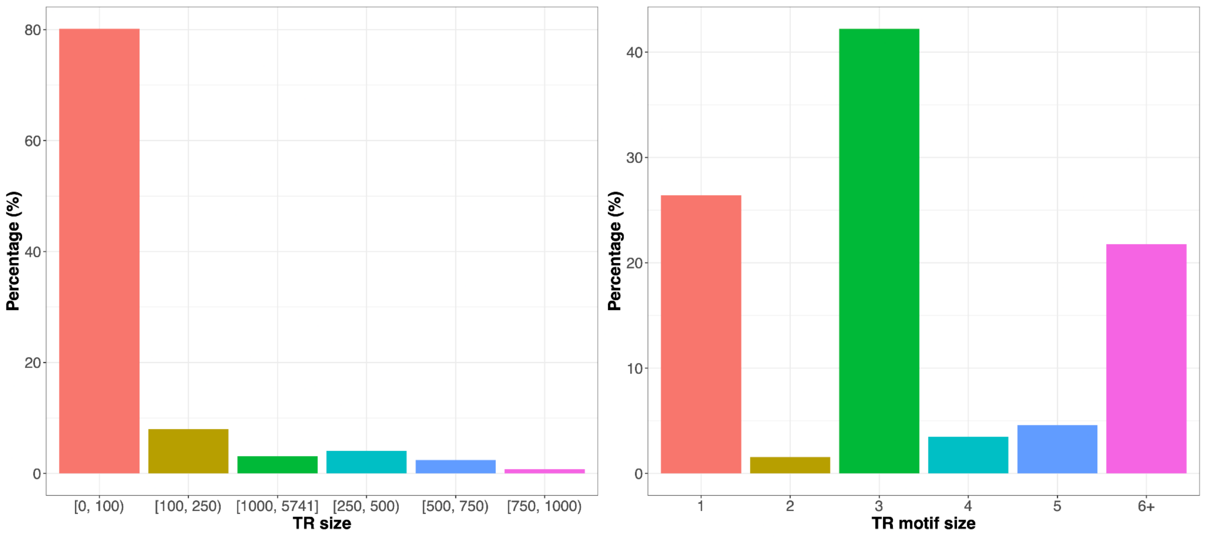


*Figure S11: Characterisation of the TRs unique to CHM13 compared to GRCh38. Left figure shows the TR allele size distribution: TR sizes were subdivided in bins. Right figure shows the distribution of TR motif sizes, as performed with TREAT/otter.*

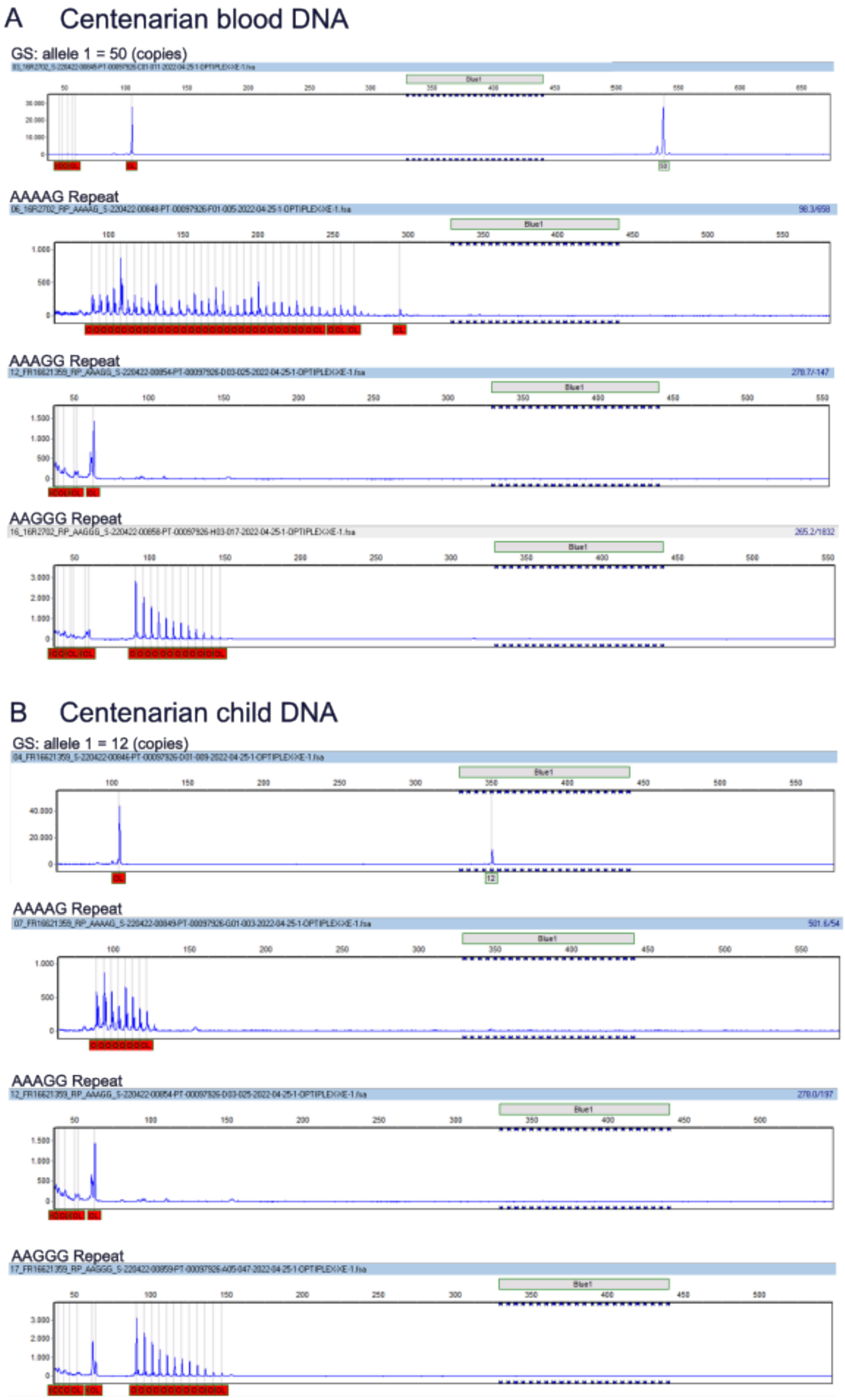


*Figure S12: Electropherogram resulting from the Repeat-Primed PCR (RP-PCR), showing experimental validation of the presence of the expanded allele in the parent as well as the*

*child.* ***A.*** *Experimental validation of the two alleles in the centenarian parent.* ***B.*** *Experimental validation of the two alleles in the centenarian child.*

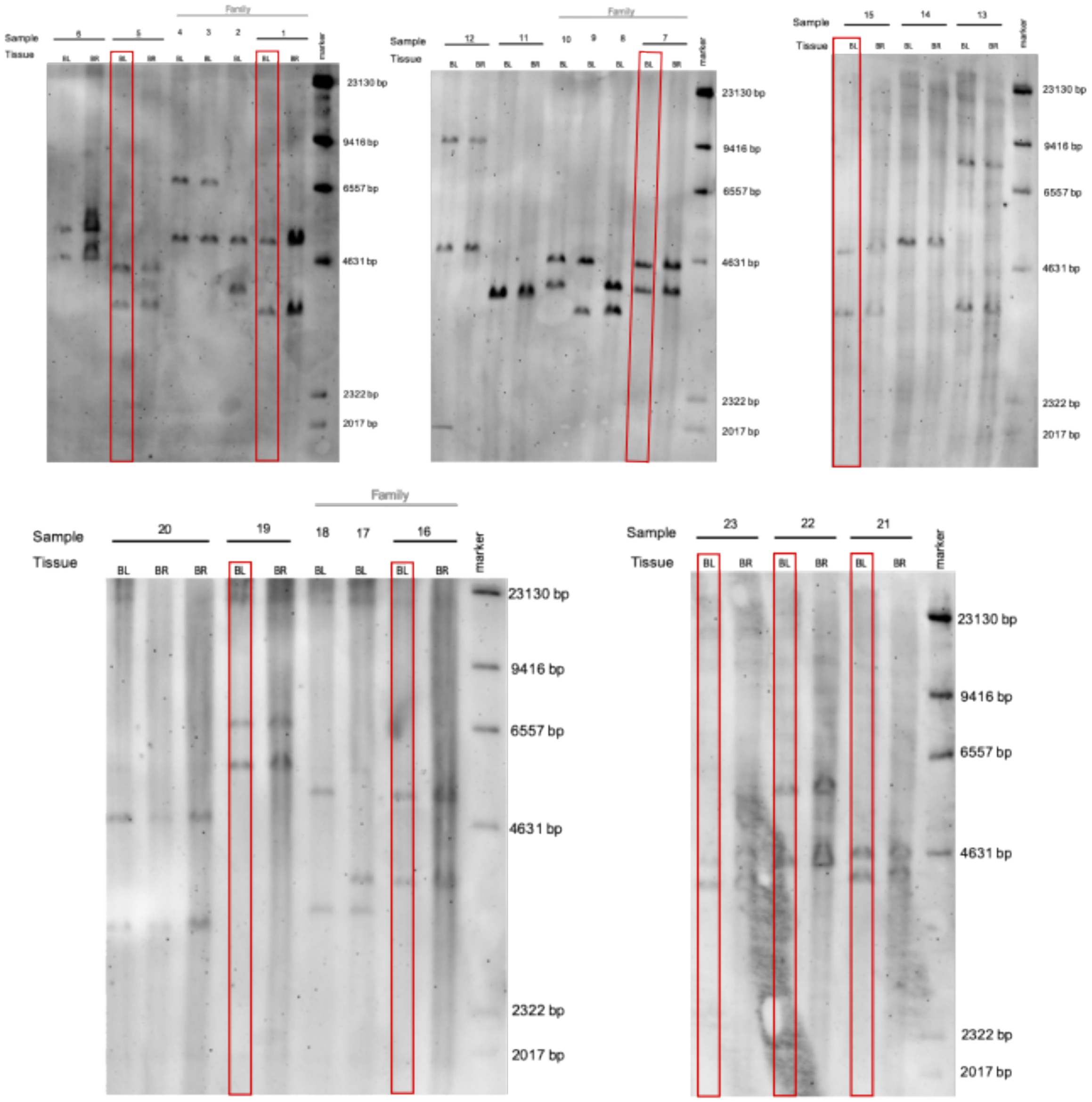


*Figure S13: Southern blot showing experimental validation of the alleles relative to the intronic ABCA7 TRs. Due to restriction digestion, flanking sequences are present around the VNTR of a size of 2038 bp. To calculate the molecular weight of the ABCA7-VNTR alleles, 2038 bp was subtracted from the size of the observed bands. Experimental validation was performed for 39 DNA samples in total. Of these, 15 were cognitively healthy centenarians (1, 5, 4, 7, 11, 12, 13, 14, 15, 16, 19, 20, 21, 22, 23). For each centenarian, we evaluated blood DNA (BL) as*

*well as brain DNA (BR) DNA. Sequencing data was available for 9/15 centenarians (blood DNA), which are highlighted in red, and these were used for the comparison of TR alleles between southern blot and sequencing-derived.*

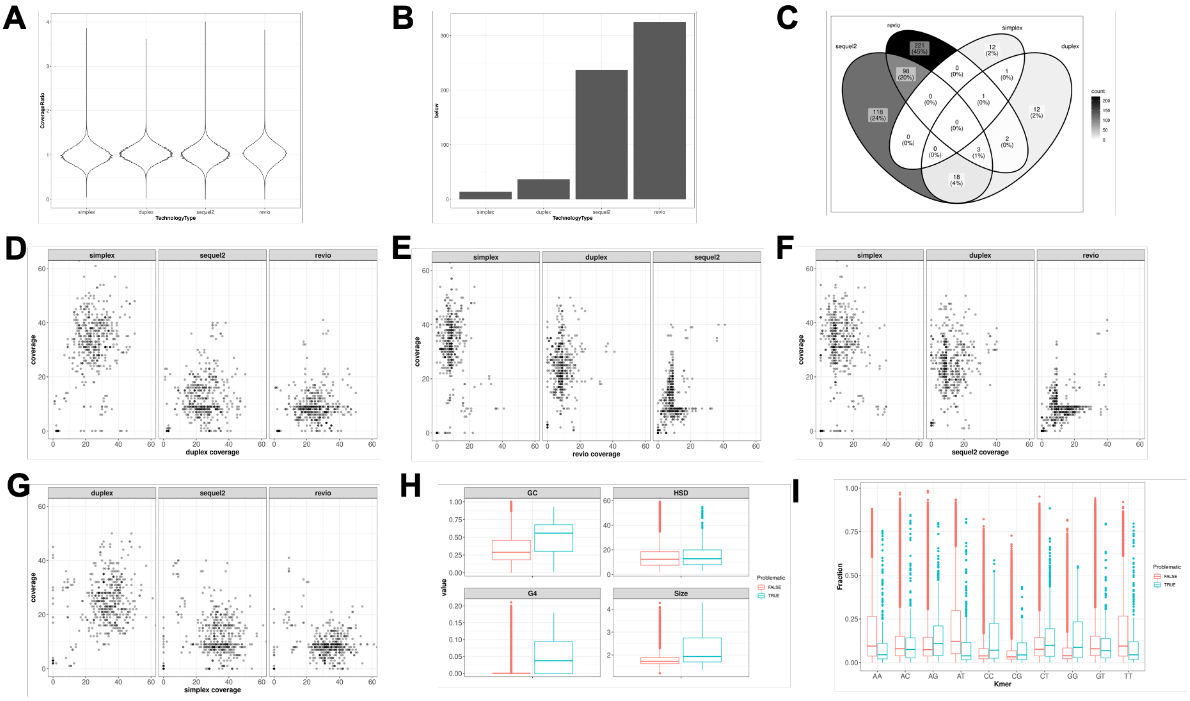


*Figure S14: Systematic analysis of coverage drops.* ***A.*** *Distribution of local vs. global ratio, for each technology.* ***B.*** *Number of problematic regions identified, for each technology.* ***C.*** *Venn showing the overlap between problematic regions across technologies.* ***D-G.*** *Shows the pairwise comparison of TR coverage across sequencing technologies in problematic regions in Duplex (D), Revio (E), Sequel 2 (F), and Simplex (G). Please note that the x-axis range was deliberately kept the same as the y-axis range.* ***H.*** *Comparison of sequence characteristics between the problematic and non-problematic regions.* ***I.*** *Dinucleotide frequency between problematic and non-problematic regions.*

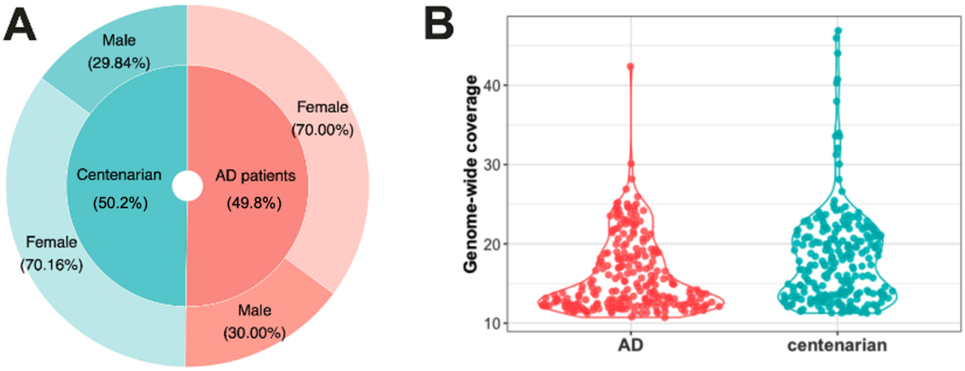


*Figure S15: PacBio HiFi sequencing of 246 Alzheimer's disease patients and 248 cognitively healthy centenarians.* **A.** Fraction of AD patients and cognitively healthy centenarians, and the relative gender characteristics. **B.** Coverage distribution of the AD patients and cognitively healthy centenarians. Individuals were sequenced with a Sequel2 instrument using 1-4 SMRT cells, for each individual.

### 3. Supplementary Methods

#### 3.1 Long-read sequencing with PacBio Sequel II and publicly available data from ONT and PacBio instruments

We downloaded publicly available unaligned long-reads from ONT 'Duplex' long-read sequencing data from the Human Pangenome Reference Consortium (HPRC). Specifically, we downloaded all available FASTQ files R10.4 chemistry base-called with 'Dorado_v0.1.1' available in the AWS HPRC bucket (https://registry.opendata.aws/hpgp-data/submissions). We aligned the long-read data to the GRCh38 (build GRCh38.p14) and the CHM13 reference genome (v2.0) using *minimap2* (2.21-r1071) with the '-x map-ont' command.[1]

Similarly, we downloaded publicly available PacBio Revio HiFi and non-HiFi data from HPRC. Specifically, we downloaded the 'hifi' and 'fail' unaligned BAM files of the 'm84039_230117_233243_s1' sequencing run available in the same AWS HPRC bucket. We then aligned both read-sets to GRCh38 and CHM13 using *pbmm2* (version 1.3.0) with the '--preset CCS --unmapped'.[2]

Finally, we generated long-read sequencing of HG002 with PacBio's Sequel 2. Briefly, raw reads are first processed using PacBio's ccs algorithm (v6.0.0, https://github.com/PacificBiosciences/ccs) to generate high-fidelity (HiFi) reads with custom parameters (min-passes 0, min-rq 0, keeping kinetics information). We retained both HIFI reads (read-quality >99%, number of passes >3), and lower-quality non-HiFi reads. We then separately merged all HiFi and non-HiFi reads, and aligned them to the GRCh38 and CHM13 reference genome using *pbmm2* using the --preset CCS --unmapped command for HIFI reads, and the --preset SUBREAD --unmapped command for non-HIFI reads.

#### 3.2 Comparison of *otter* and TREAT with existing tools

*Generation of data with reduced coverage*: For each of the four long-read datasets (PacBio Sequel 2 and Revio, and ONT Simplex and Duplex), we generated six reduced-coverage BAM files using *samtools* (v1.13) '*view'* command with the '-s' argument to randomly reduce genome-wide coverage to 5x, 10x, 15x, 20x, 25x, and 30x.[1] This was done for both GRCh38 and CHM13 aligned datasets.[3]

*Generation of ground truth dataset of HG002*: To generate the 'ground-truth' set of the expected allele-sequences for the 161K TRs in HG002, we whole-genome aligned the HG002 T2T assembly (v0.7, https://github.com/marbl/HG002) to GRCH38 using *minimap2* with the '-*ax asm10*' parameter.[2] We extracted the corresponding allele sequences per parental and maternal contigs by iterating through each contig-alignment and extracting the corresponding TR sequences by calculating the start and end coordinates using the CIGAR string. We omitted TRs that were non-spanning (*e.g.* soft/hard-hard clipped alignments within the TR sequence), originated from unexpected chromosomes (*e.g.* mismatching chromosomes between GRCH38 and CHM13), and had more or less than two alignments (*e.g.* two parental sequences). We were able to identify a total of 322,772 parental and maternal allele-sequences in the HG002 T2T assembly, corresponding to 161,111/161,382 (99.8%) GRCh38-annotated TRs.

*Curated set of 864K TRs*: We downloaded the CHM13 reference genome as well as the following repeat annotations from the UCSC Genome Browser (https://hgdownload.soe.ucsc.edu/gbdb/hs1/): *SimpleRepeat*, *RepeatMasker*, *CenSat*, and *SegDups*. We extracted *Satellite* and *Simple_repeat* annotations from *RepeatMasker*,[3] and combined them with annotations from SimpleRepeat, merging all annotations within <=100 bp of each other. We excluded annotations that were <=10 Kbp of centromeric sequences

(Censat) and segmental duplications (SegDups), and removed TRs from both sex chromosomes.

*Evaluation of genotyping accuracy*: We evaluated genotype accuracy by comparing the reported TR allele-sequences of each tool with the expected allele-sequences from the HG002 T2T assembly. Briefly, we assume at most two observed and expected allele sequences per TR, and duplicate single observed allele-sequences, as some tools (*e.g. otter*) reports only a single allele-sequence per TR in homozygous situations. Note that this differs from TRs that are deleted, which are designed as 'N' and 'NDNNN' sequences in *otter* and LongTR, respectively. For each tool and each TR, we calculated two distances using the following procedure: let {o1, o2} and {e1, e2} be the observed and expected allele-sequences for a given TR, and *dist(x, y)* be the edit distance for strings *x* and *y*. We compute dist(o1, e1), dist(o1, e2), dist(o2, e1), dist(o2,e2), and used the two distances with the minimum values. The raw edit distance was calculated using WFA-lib2 (https://github.com/smarco/WFA2-lib).[4] The normalised edit distance was obtained as the ratio between the raw edit distance, and the size of the expected allele.

*Selection of genes from CHM13 reference genome*: The full list of genes in CHM13 was downloaded from UCSC Table Browser (https://genome.ucsc.edu/cgi-bin/hgTables), specifying *T2T CHM13v2.0/hs1* as Assembly, *NCBI RefSeq* as Group under *Genes and Gene Predictions*, and *RefSeq All (hub_3671779_ncbiRefSeq)* Table. This table was further curated by removing *antisense* transcript, Long Intergenic Non-Coding RNAs (LINC), and RNA genes. In case multiple occurrences of the same gene existed, the largest gene (based on *chromStart* and *chromEnd* coordinates) was used. The curated list of genes included 24,429 entries.

*Commands used for benchmarking*: Genotyping accuracy was based on running *otter* standalone with the following command:

```
otter assemble -A 5000,0.1 -c 150 -b [input.bed] -r [reference.fasta]
-R [output_name] -t 4 [input.bam] | samtools view -bh | samtools sort
> [output_name].bam
```

Motif accuracy and running performances was based on running TREAT's unified workflow:

```
TREAT assembly -i [input.bam] -b [input.bed] -r [reference.fasta] -t
4 -o [output_dir] -w 20 -wAss 20
```

The command used to evaluate genotyping and motif accuracy in TRGT was the following:[5]

```
TRGT genotype -g [reference.fasta] -r [input.bam] -b [input.bed] -o
[output_name] -t 4
```

The command used to evaluate genotyping accuracy in LongTR was the following:[6]

```
LongTR –bams [input.bam] –fasta [reference.fasta] –regions
[input.bed] –tr-vcf [output_name].vcf.gz –skip-assembly –bam-samps
[sample_name] –bam-libs [sample_name] –max-tr-len 5000 –min-reads 5
```

#### 3.3 TREAT

*Reads analysis:* The *reads* analysis extracts all reads spanning a given region of interest using Samtools.[3] Then, TR genotyping is performed using an iterative clustering framework. A minimum number of reads supporting each allele needs to be satisfied (by default, this equals 2). An iterative k-means clustering (k=2) framework is used to identify the TR allele sizes, while controlling for a maximum deviation of the TR sizes within each allele based on the median absolute deviation (MAD, by default, 10% of median allele size, modifiable with the '--haploDev' parameter). This framework recognizes homozygous calls and its iterative nature makes it robust to outliers. TR content characterization (motif and number of copies) in the *reads* analysis first identifies motifs and copy numbers at the individual read level (see *Methods*). Then, the most frequently observed motifs across all individual reads are prioritised.

The output VCF file produced with the *reads* analysis does not contain the allele sequences, but only allele size and motif composition.

*Motif identification, identical both assembly and reads analyses*: For motif content characterization, *pytrf* python extension is used (https://github.com/lmdu/pytrf). First, we run pytrf with default parameters (specifying min_motif_size=1 and max_motif_size=100). In case no repeat pattern is found, we relax pytrf parameters (min_seed_repeat=2 and min_seed_length=8).

*Motif representation:* TREAT reports two representations of TR motifs and relative copy numbers: an individual-based and a reference-based. The individual-based motif represents the motif observed in an individual's TR alleles. The reference-based motif forces the copy number estimation to be based on the size of the motif found in the reference genome. Assuming a TR with the sequence "AGGA AGGA CGGC CGGC" in an individual, with the relative TR in the reference genome being "AGGA AGGA AGGA", (*i*) the reference-based motif would be *AGGA* (the motif observed in the reference genome), repeated 4 times in the individual and 3 time in the reference genome, and (*ii*) the individual-specific representation would be *(AGGA)2(CGGC)2*. Both motifs are generated and reported to the user in the gVCF/VCF file.

*Outlier analysis*: In the outlier analysis, TREAT optionally accepts a file reporting TR-specific pathogenic thresholds, and will report any instance of TRs exceeding the provided threshold in any individual.

*TREAT commands used in this study*: TREAT *assembly* analysis was done using the following command:

```
TREAT.py assembly -b input.bed -i input1.bam,input2.bam,input3.bam -
o outputDirectory -w 20 -wAss 20 -t 4 -r reference_ghcr38.fa
```

The above command was used (*i*) in the comparison of alleles between the *reads* and *assembly* analyses; (*ii*) in the analysis of clinically relevant regions using the genomes of 47 HPRC, the 2 CANVAS, and the 10 parent-child duos; (*iii*) for the benchmarking with other tools in the evaluation of motif identification accuracy; (*iv*) in the case-control analysis.

The *reads* analysis was performed using the following command:

```
TREAT.py reads -b input.bed -i input1.bam,input2.bam,input3.bam -w 20
-t 4 -o outputDirectory -r reference_grch38.fa
```

The above command was used (*i*) in the comparison of alleles between the *reads* and *assembly* analyses.

TREAT plots were generated with the following command:

```
TREAT.py plots -v samples.vcf.gz -o plots_folder -r all
```

TREAT outlier analysis was performed with the following command:

```
TREAT.py analysis -a outlier -v samples.vcf.gz -o analysis_folder -r
all
```

TREAT case-control analysis was performed with the following command:

```
TREAT.py analysis -a case-control -v samples.vcf.gz -l
case_control_labels.txt -o analysis_folder -r all
```

#### 3.4 *Otter*

*Otter algorithm details*: In the context of TR genotyping, *otter* will attempt to identify a candidate set of unique allele-sequences by clustering spanning-reads via pairwise-sequence alignment. To deal with high somatic variation and/or sequencing errors, *otter* will first estimate local base-line error-rates per region. More specifically, we apply a gaussian-kernel density estimator across a one-dimensional distribution of all spanning pairwise-sequence distances in a region. For single homozygous allele-sequences, this distribution will be unimodal with centre at 0, representing the baseline error-rate of the sequences. When two or more unique allele-sequences are present, the distribution will be multi-modal such, that ε0, ε1, ε2, …, εm are global maximas, and ε1, ε2, …, εm represent sequence errors between reads from different allele-sequences. We thus identify global maximas and minimas through a simple peak-finding procedure by sampling n points (by default N=100) through this distribution. *Otter* uses this information to perform a hierarchical clustering of the reads and stopping when the distance exceeds the densest peak among ε1, ε2, …, εm, ultimately partitioning reads into initial clusters representing candidate allele-sequences.

Non-spanning reads are assigned to each cluster by aligning each non-spanning read to all spanning reads and identifying the closest sequence based on maximum sequence similarity. Cluster membership of closest spanning read is extended to the non-spanning read if it meets a given sequence similarity threshold (default 90%), and if the sequences similarity of spanning reads in other clusters differs by no less than a given threshold (default 1%).

*Otter* then generates consensus sequences per cluster based on a pseudo-partial order alignment procedure inspired by *Ye and Ma, 2016.*[7] First, *otter* identifies a representative sequence per cluster by identifying the spanning read that minimises the global sum of pairwise distances. This read, regarded as the backbone of the consensus, is then converted to a weighted directed acyclic graph (DAG) where each nucleotide is an individual node, and the weight of each edge represents the number reads traversing two pair of nodes (initially set to 1). All other reads in the cluster (both spanning and non-spanning) are aligned to the representative read via an affine-gap alignment, and the DAG is updated with new nodes, edges, and weights. The weights are normalised based on scaling procedure described by *Ye and Ma, 2016*,[7] and the heaviest weighted path in the DAG is identified by maximising the path with total sum of weighted edges, solved via dynamic programming. The heaviest weighted path is then chosen as the consensus sequence per cluster.

*Otter commands used in this study*: Benchmarking of *otter* across 864K TRs in ONT's Duplex and Simplex data, as well as PacBio's Revio an Sequel 2 data was performed with four threads, a coordinate offset of 5 nucleotides, and local realignment with the (given) CHM13 reference genome via the following command:

```
otter assemble -t 4 -r chm13.fa -o 5 --sam -R sample sample.bam
```

Integration of non-HIFI data for the assembly of intronic TRs in *RFC1* and *ABCA7* was performed with the following additional parameters:

```
-s 0.8 -f 200 -A500,0.1
```

This enables more sensitivity during local re-alignment: realign 200-bp flanking sequence with a minimum sequence similarity of 80%, and coverage thresholds for clusters containing allele-sequences above 500 bp to 10%.

#### 3.5 Cohort and sequencing details

*Cohort of Alzheimer's Disease patients and cognitively healthy centenarians:* We whole-genome sequenced using PacBio Sequel 2 instrument 494 individuals. Of these, 246 were clinically diagnosed with probable AD from the Amsterdam Dementia Cohort (ADC).[8] The diagnosis of probable AD in ADC cohort was based on the clinical criteria formulated by the National Institute of Neurological and Communicative Disorders and Stroke – Alzheimer's Disease and Related Disorders Association (NINCDS-ADRDA) and based on the National Institute of Aging-Alzheimer Association (NIA-AA). All subjects underwent a standard diagnostic assessment including neurological examination, blood tests, magnetic resonance imaging, electroencephalogram, and cerebrospinal fluid (CSF) analysis (available for most patients). In addition to AD patients, we sequenced 248 cognitively healthy centenarians from the 100-plus Study.[9] This study includes Dutch-speaking individuals who (*i*) can provide official evidence for being aged 100 years or older, (*ii*) self-report to be cognitively healthy, which is confirmed by a proxy, (*iii*) consent to the donation of a blood sample, (iv) consent to (at least) two home visits from a researcher including an interview and neuropsychological test battery. The Medical Ethics Committee of the Amsterdam UMC approved all studies. All participants and/or their legal representatives provided written informed consent for participation in clinical and genetic studies.

*Sequencing with PacBio Sequel 2*: All samples were sequenced using a PacBio Sequel 2 instrument. Specific details about the sample preparation, sequencing, and data analysis, are available elsewhere.[9] Briefly, raw reads are first processed using PacBio's ccs algorithm (v6.0.0, https://github.com/PacificBiosciences/ccs) to generate high-fidelity (HIFI) reads with custom parameters (min-passes 0, min-rq 0, keeping kinetics information).[10] We retained both HiFi reads (read-quality >99%, number of passes >3), and lower-quality non-HiFi reads. We then separately merged all HiFi and non-HiFi reads, and aligned them to the GRCh38 and CHM13 reference genome using *pbmm2* using the '--preset CCS --unmapped' command for HiFi reads, and the '--preset SUBREAD --unmapped' command for non-HiFi reads.[10]

#### 3.6 Southern Blotting

Southern blotting (SB) assay was used to determine the size of the TR in *ABCA7* gene (chr19:1049514-1049953, based on GRCh37) for 11 cognitively healthy centenarians from the 100-plus Study. gDNA from blood samples were isolated using automated isolation technology (PerkinElmer Chemagen Technology) and stored at 4°C. To check if isolated gDNA was high in molecular weight it was run on a 0.8% agarose gel containing ethidium bromide on 220 volts for 60-70 minutes. Hereafter 3 µg of gDNA isolated from blood material underwent restriction digestion for 5-6 hours at 37°C using BamHi (New England Biolabs) according to manufactures' protocol.

*Gel-electrophoresis and blotting*: Samples that underwent restriction digestion were run on a 0.8% agarose gel (agarose LE, Lonza) for ~17 hours overnight (O/N) on 60V. DIG molecular weight marker #2 (Roche) was used as a marker to determine size of the DIG-labelled DNA fragment of interest later on. After running, to check for full digestion of the gDNA, gel was incubated in MilliQ-water containing ~0.2-0.5 µg/mL ethidium bromide for 30 minutes, whereafter DNA was visualised using UV-light. Gel was then rinsed with MilliQ-water (MQ-water) and incubated in a depurination buffer (250mM HCl) for 10 minutes, gently shaking at room temperature (RT). Hereafter, gel was rinsed with MQ-water and incubated in a denaturation buffer (0.5 M NaOH; 1.5 M NaCl) for 30 minutes, gently shaking at RT. Gel was then rinsed with MQ-water and incubated in a neutralisation buffer (0.5 M Tris-HCl pH 7.5; 1.5 M NaCl) for 30 minutes, gently shaking at RT. After this, gel was equilibrated in a 20x SSC buffer for a minimum of 10 minutes whilst building the blotting set up (description of full set up can be found in appendix 1). Blotting of the DNA onto a positively charged nylon membrane (Roche) was carried out.

*Hybridization of DIG-labelled probe*: After blotting, the membrane was retrieved from the blotting set-up, whereafter the DNA on the membrane was crosslinked for 3 minutes using UV-light. The membrane was then rinsed using MQ- water and transferred into a hybridization bottle where it was incubated in preheated (52°C) DIG Easy Hyb (Roche) for 2 hours, rotating at 52°C. DIG labelled probe was generated using PCR DIG Probe Synthesis Kit (Roche). Primers were designed as described by deRoeck et al., 2018.[11] The following primers were used: forward primer: 5'-AGCTCTGTAACTGCCAGTGC-3'; reverse primer: 5'-CCGTAGGCTCGTCCAGGAT-3'. Probe was denatured by cooking at 95°C for 5 minutes, immediately chilling on ice afterwards. To make a hybridization solution, preheated (52°C) DIG Easy Hyb (Roche) was combined with a denatured probe. Membrane was incubated in hybridization solution 16 hours O/N, rotating at 52°C.

*Imaging*: After hybridization, the membrane was incubated in a low stringency buffer (2x SSC containing 0.1% SDS) for 5 minutes, gently shaking at RT, then rinsed with MQ water and incubated again in low stringency buffer for 5 minutes gently shaking at RT. Hereafter, membrane was incubated in preheated (68°C) high stringency buffer (0.1x SSC containing 0.1% SDS) for 15 minutes, gently shaking at 68 °C and rinsed with MQ water thereafter incubating in preheated high stringency buffer again for 15 minutes, gently shaking at 68°C. Then, the membrane was incubated in a washing buffer (made fresh using DIG Wash and Block Buffer Set (Roche)) for 2 minutes gently shaking at RT. After washing, the membrane was incubated in a freshly made blocking buffer (DIG Wash and Block Buffer Set (Roche)) for 1.5 – 2.5 hours shaking gently at RT. Antibody solution was made using Anti-Digoxigenin-AP antibody (Roche) diluted to 1:10000 in blocking buffer. Membrane was incubated in antibody solution for 30 minutes, shaking gently at RT. Hereafter, the membrane was washed twice for 15 minutes using a washing buffer, gently shaking at RT. Membrane was then equilibrated in a freshly made detection buffer (DIG Wash and Block Buffer Set (Roche)) for 3 minutes, shaking gently at RT. Membrane was placed on a transparent sheet thereafter CDP-star substrate (Roche) was applied dropwise to the membrane, immediately covering applied areas with a second transparent sheet to prevent dehydration. Membrane was then protected from the light when transferring to the imager (UVITEC Cambridge Alliance 4.7). Imaging was performed for 10 to 15 minutes using the chemiluminescent option on the imager, whereafter images were analysed using the Alliance software provided by UVITEC.

*Analysis*: Using the alliance software, the molecular weight of the band signals on the membrane representing ABCA7-VNTR allele length were determined by comparison with known marker fragment sizes. Due to restriction digestion flanking sequences are present around the VNTR of a size of 2038 bp. To calculate the molecular weight of the ABCA7-VNTR in the analysed samples, 2038 bp was subtracted from the size of the observed bands.

#### 3.7 A curated set of genome-wide set TRs in CHM13

We curated a genome-side set of TR in CHM13 based on available annotations at UCSC genome browser (downloaded March 3, 2023). First, we downloaded all available repeat annotations from RepeatMasker and retained all available TR-annotations. We then remove TRs that overlapped or were within 10 Kbp of centromeric and/or segmental duplications (>= 5 Kbp) annotations. All resulting TRs within 100 bp of each other were merged to a single composite TR annotation, and TRs > 5 Kbp were removed. This forms the curated set of genome-wide TRs in CHM13. For CHM13-unique TRs, we used UCSC *liftOver* (version downloaded March 1, 2023) to 'lift' the TR annotations to the GRCH38 reference genome, and retained only those that were fully 'Deleted' as reported by *liftOver*.

### 5. References

1. Li, H. Minimap2: pairwise alignment for nucleotide sequences. *Bioinformatics* **34**, 3094–3100 (2018).
2. Wenger, A. M. *et al.* Accurate circular consensus long-read sequencing improves variant detection and assembly of a human genome. *Nat Biotechnol* **37**, 1155–1162 (2019).
3. Danecek, P. *et al.* Twelve years of SAMtools and BCFtools. *Gigascience* **10**, giab008 (2021).
4. Marco-Sola, S. *et al.* Optimal gap-affine alignment in *O* ( *s* ) space. *Bioinformatics* **39**, btad074 (2023).
5. Dolzhenko, E. *et al.* Characterization and visualization of tandem repeats at genome scale. *Nat Biotechnol* (2024) doi:10.1038/s41587-023-02057-3.
6. Ziaei Jam, H. *et al.* LongTR: genome-wide profiling of genetic variation at tandem repeats from long reads. *Genome Biol* **25**, 176 (2024).
7. Ye, C. & Ma, Z. (Sam). Sparc: a sparsity-based consensus algorithm for long erroneous sequencing reads. *PeerJ* **4**, e2016 (2016).
8. van der Flier, W. M. & Scheltens, P. Amsterdam Dementia Cohort: Performing Research to Optimize Care. *Journal of Alzheimer's Disease* **62**, 1091–1111 (2018).
9. Holstege, H. *et al.* The 100-plus Study of cognitively healthy centenarians: rationale, design and cohort description. *European Journal of Epidemiology* (2018) doi:10.1007/s10654-018-0451-3.
10. Salazar, A. *et al. An AluYb8 Retrotransposon Characterises a Risk Haplotype of TMEM106B Associated in Neurodegeneration*. http://medrxiv.org/lookup/doi/10.1101/2023.07.16.23292721 (2023) doi:10.1101/2023.07.16.23292721.
11. On Behalf of the BELNEU Consortium *et al.* An intronic VNTR affects splicing of ABCA7 and increases risk of Alzheimer's disease. *Acta Neuropathologica* **135**, 827–837 (2018).
